# Transient dopamine neuron activity precedes and encodes the vigor of contralateral movements

**DOI:** 10.1101/2021.04.20.440527

**Authors:** Marcelo D Mendonça, Joaquim Alves da Silva, Ledia F. Hernandez, Ivan Castela, José Obeso, Rui M Costa

## Abstract

Dopamine neurons (DANs) in the substantia nigra pars compacta (SNc) have been related to movement vigor, and loss of these neurons leads to bradykinesia in Parkinson’s disease. However, it remains unclear whether DANs encode a general motivation signal or modulate movement kinematics. We imaged activity of SNc DANs in mice trained in a novel operant task which relies on individual forelimb sequences. We uncovered that a similar proportion of SNc DANs increased their activity before ipsi- *vs*. contralateral sequences. However, the magnitude of this activity was higher for contralateral actions, and was related to contralateral but not ipsilateral sequence length. In contrast, the activity of reward-related DANs, largely distinct from those modulated by movement, was not lateralized. Finally, unilateral dopamine depletion impaired contralateral, but not ipsilateral, sequence length. These results indicate that movement-initiation DANs encode more than a general motivation signal, and invigorate kinematic aspects of contralateral movements.

**Teaser:** Transient activity in substantia nigra compacta dopamine neurons encodes contralateral, but not ipsilateral action vigor.

## Introduction

Choosing which actions to perform in specific contexts is critical for survival. It is also critical to perform these actions at the right time and with the right potency, i.e. force, speed, duration. Basal ganglia circuits, and dopaminergic signaling in these circuits, are critical for the modulation of both movement initiation and movement vigor (1, 2). Accordingly, one essential feature of Parkinson’s Disease (PD), which is characterized by a progressive loss of dopaminergic neurons (DANs) in the Substantia Nigra *pars compacta* (SNc) (3), is reduced amplitude (hypokinesia) and slowness of movements (bradykinesia).

Early studies identified that substantia nigra *pars compacta* (SNc) dopaminergic activity was modulated during large reaching movements (4, 5). More recently, the activity of DANs was found to be transiently modulated around movement onset (1, 2, 6–9), and manipulations of this activity before movement onset had an impact on the probability of movement execution and the vigor of movements (1, 2). This is supported by well-established observations that chronic DA depletion leads to decreased amplitude, peak force, and speed of movement in PD (10–12), and also in rodents (13–15). While many studies of vigor and PD have focused on movement force and speed, the length/duration of movement sequences is also critically affected, and has been less studied. For example, gait bouts of PD patients are characterized not only by a lower speed but also by a reduced number of steps per bout (16).

It has been proposed that DA neurons influence movement vigor by modulating the motivation to behave (17, 18). Behavioral studies in PD revealed that the deficit in movement vigor reflects a reduced probability of committing to more vigorous actions, even when necessary for obtaining a reward (12). PD motor signs typically start focally on one side of the body (19), contralateral to the most denervated SNc, where movement vigor deficits are observed (20). Similarly, unilateral dopamine depletion in mice leads to deficits in contraversive, but not ipsiversive movements (21). Striatum is involved in contralateral movements (22–24), and DANs activity is higher when animals perform contralateral versus ipsilateral choices (7). Thus, dopaminergic activity is properly placed to affect movement kinematics in a lateralized way by directly influencing medium spiny neurons (MSNs) excitability (25).

In this study, we investigate the hypothesis that movement-related DANs signal not only a general motivation to move, but invigorate specific kinematic aspects of contralateral movements. Towards this end, we developed a novel behavioral task where freely moving mice have to perform fast movement sequences using an individual forelimb in order to obtain reward. This paradigm allowed us to investigate movement sequences performed with either forelimb. We imaged the activity of genetically identified SNc DA neurons using one-photon imaging during lever press task performance. We identified distinct populations of SNc DANs with transient activity related to movement *versus* reward. Although a similar proportion of movement-related neurons were observed bilaterally, their activity was higher for contralaterally performed sequences than for ipsilateral ones. Furthermore, this movement-related activity was related to sequence length but only for contralateral sequences. In contrast, reward-related activity was not lateralized. Consistently, unilateral lesion of SNc dopaminergic neurons led to contralateral, but not ipsilateral, forelimb vigor impairment. These results suggest a role of transient dopaminergic activity before movement in invigorating the duration of contralateral movements.

## Results

### Mice learn to perform rapid single-forelimb lever press sequences

We trained mice (n=8) to perform a fast lever-pressing task where it was required to press a lever at increasingly higher speed in order to obtain a 10% sucrose reward. During the training, spatial constraints in the lever were imposed in order to restrict the accessibility of the lever forcing the animal to use only one specific forelimb (Fig 1A, Movie S1, Movie S2).

**Figure 1:**
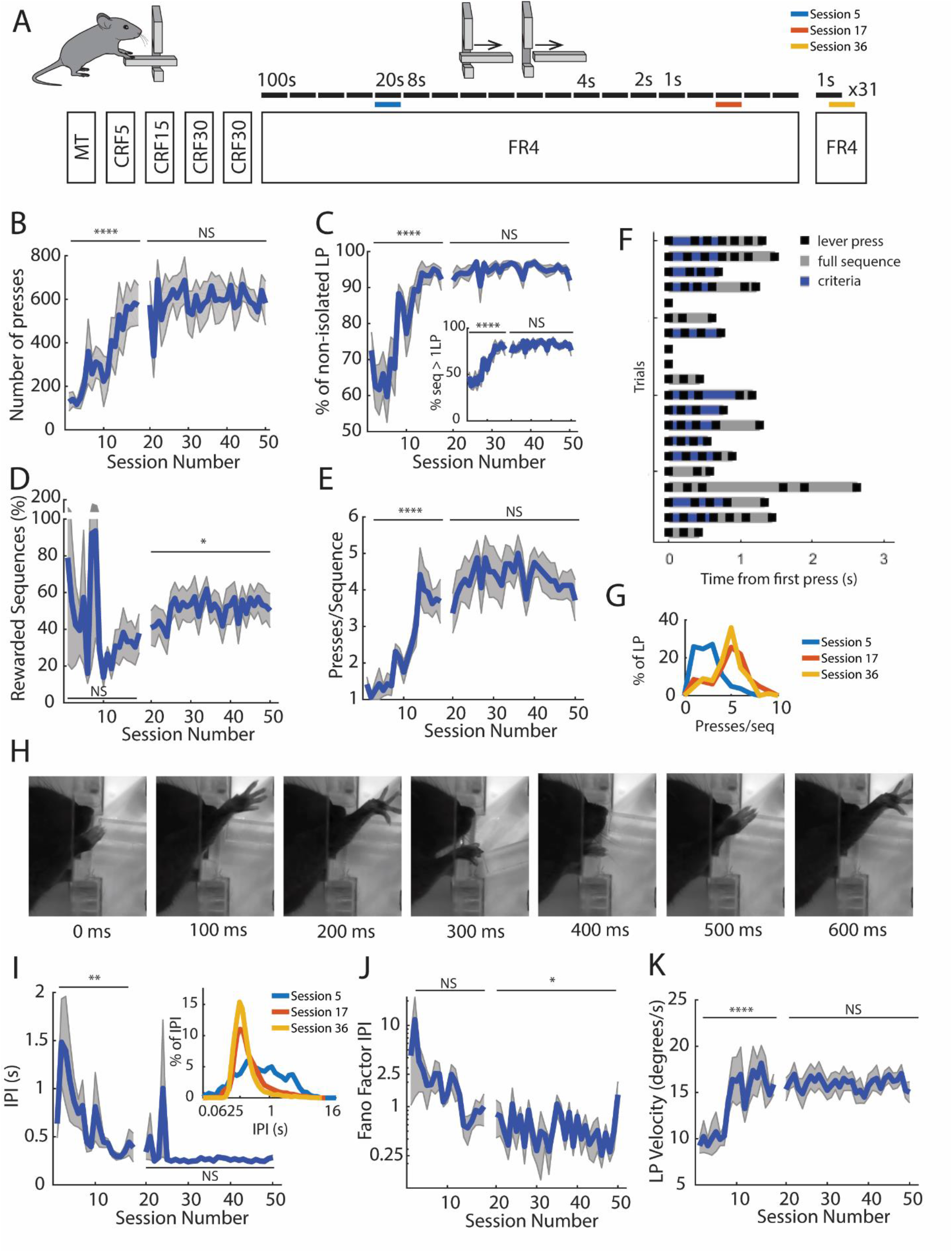
A novel task for assessment rapid single-forelimb lever press sequences. **A)** Schematics of the training schedule. Training starts with a first session of Magazine Training (MT) followed by 4 sessions of continuous reinforcement schedule (CRF) where each press leads to one reward. Animals were trained for 19 sessions with an FR4 schedule of increasing time constraint (moving from 4 presses in 100 seconds to 4 presses in 1 second). During this period, there was also the progressive retraction of the lever that led to lever presses (LP) performance with a single forelimb. After this period, Animals were moved to a performance phase during 31 sessions, where 4 lever presses performed in less than one second led to a reward. The blue, orange and yellow lines represent example sessions to be specifically illustrated in Figures 1G and 1I. In panels B), C), D), E), J) and K), there is a discontinuation in the line. This represents the end of the stage defined as training, and the beginning of the stage defined as performance. **B)** With training, the total number of Lever Presses increased and **C)** animals rapidly started to organize their behavior in self-paced bouts or sequences of lever presses, until there were almost no single presses occurring. **D)** In the performance stage about 60% of sequences were rewarded as animals reorganize their behaviour **G)** and start to perform lever press sequences of 4-5 presses **E)**. **F)** Example of sequences performed by a representative animal, aligned at the time of sequence initiation. Individual lever presses are marked as black ticks, the full sequence duration is shaded in grey and the IPIs that meet the session minimum target are shaded in blue. **H)** Representative frames collected from a high-speed (120 fps) camera during sequence performance. **I)** With training mice decrease the Inter-press interval to a mean of 0.347 seconds. The reorganization of IPIs distribution (*Inset*) happens during training. **J)** Variability of the inter-press interval decreases while press velocity increases across training **K)**. (Error bar denotes S.E.M.) *p<0.05; **p<0.01; ***p<0.001; **** p<0.0001. For detailed statistical analysis, see Table S1.

After introducing the animals to the apparatus and 4 days of continuous reinforcement (CRF, one press = one reward), animals were trained at a progressively faster fixed-ratio schedule (FR4, 4 presses = one reward), up to a maximum of 4 presses in less than 1 second. During the FR4 training, the lever was progressively receded to guarantee that it was only accessible to one forelimb (Fig 1A,H, details in the Methods section). Animals were moved across a training schedule of 19 sessions starting with FR4 in 100 seconds and ending with 5 sessions of FR4 in 1 second. Stability of this asymptotic performance, was tested in 31 consecutive FR4/1sec sessions.

With training, the increase in the total number of lever presses (Fig 1B, F(18,126) = 4,536 p<0.0001), paired the increase in the number of presses/minute (reaching 14.95 +- 12.26 presses/minute in the last session, F(18,126) = 4,254, p<0.001 Fig S1A). During this period, animals rapidly started to organize their behavior in self-paced bouts or sequences of lever presses (Fig 1C-G) with the percentage of lever presses performed within a sequence increasing significantly across training (F(18,109) = 7.526, p<0.0001; Fig 1C) until there were almost no single presses occurring in isolation. In the last training session 92.26% +- 8.62 of lever presses occurred within a sequence with 77.12% +- 19.20 of the sequences being composed by more than one LP (Fig 1C inset).

The number of lever presses within a sequence progressively increased (F(18,109) = 10.52, p<0.0001, Fig 1E,G), with the distribution of lever presses per sequence exhibiting a clear peak at 4 lever presses matching the imposed rule (3.69 +- 1.71, t7=0.5169, p=0.6212). Also, as time criteria become more demanding, mice decrease their inter-press intervals (IPIs) up to 0.387 +- 0.421 s (not significantly different from a target of 0.333, t7=0.3619, p=0.7281 an IPI corresponding to the performance of 4 presses in less than 1 second, F(18,108) = 2.134, p=0.0089, Fig 1I). Consistent with the reduction in the IPI, mice also increased their mean lever press velocity with training (Fig 1K). There was also reduction of the variability of the IPIs (Fano Factor, Fig 1J, F(49,315) = 1.115, p=0.2873, post-hoc test for linear trend: F(1,315)=16.08, p<0.0001).

After the 19 sessions of training, animals were assessed in 31 additional sessions with the criteria of 4 LPs in less than one second, after behavior had asymptote, with no substantial differences noted across these sessions in behavior metrics (Fig 1, Fig S1, Table S1).

These data indicate that animals learned to shape their behavior to get closer to the target criteria and, after learning, this performance is stable across time. Additionally, animals can be trained to use individual forelimbs (Fig S2), allowing us to test ipsi and contralaterally performed movement sequences.

### Transient activity of SNc dopaminergic neurons precedes movement sequence initiation

In order to investigate the activity of SNc DANs during the execution of contralateral vs ipsilateral movements we chronically-implanted gradient index (GRIN) lenses above the SNc (Fig S9D for lens location), and injected a genetically encoded calcium indicator (GCaMP6f) into genetically-identified dopaminergic SNc cells (DAT-Cre), and imaged the activity using a one-photon miniaturized epifluorescence microscope (26), (Fig 2a-b). Half of the animals (total n=6 mice) had a virus injection and lens implanted in the left hemisphere and the other half in the right one. We trained these animals in the same task described above, but with 2 independent sessions each day, on each session each animal had to use a distinct forelimb (Fig 2D). Each day mice were placed in a box with a lever available on one of the sides (left or right). The session ended after the animal obtained 30 rewards or 30 minutes have passed. After the first session, the animals were removed from this box, and placed in their homecage for a period of 30 to 150 minutes before being trained in a second session in the box, but with the other lever available. The order of limb trained (left or right) was pseudo-randomized across days. Mice were able to perform movement sequences with both forelimbs and, after training, no significant differences in behavioral metrics were noted between the two forelimbs (data summarized in Fig S3, Table S2, and presented as ipsilateral and contralateral forelimb to the implanted lens). Neural activity was assessed during the phase of asymptotic performance.

**Figure 2:**
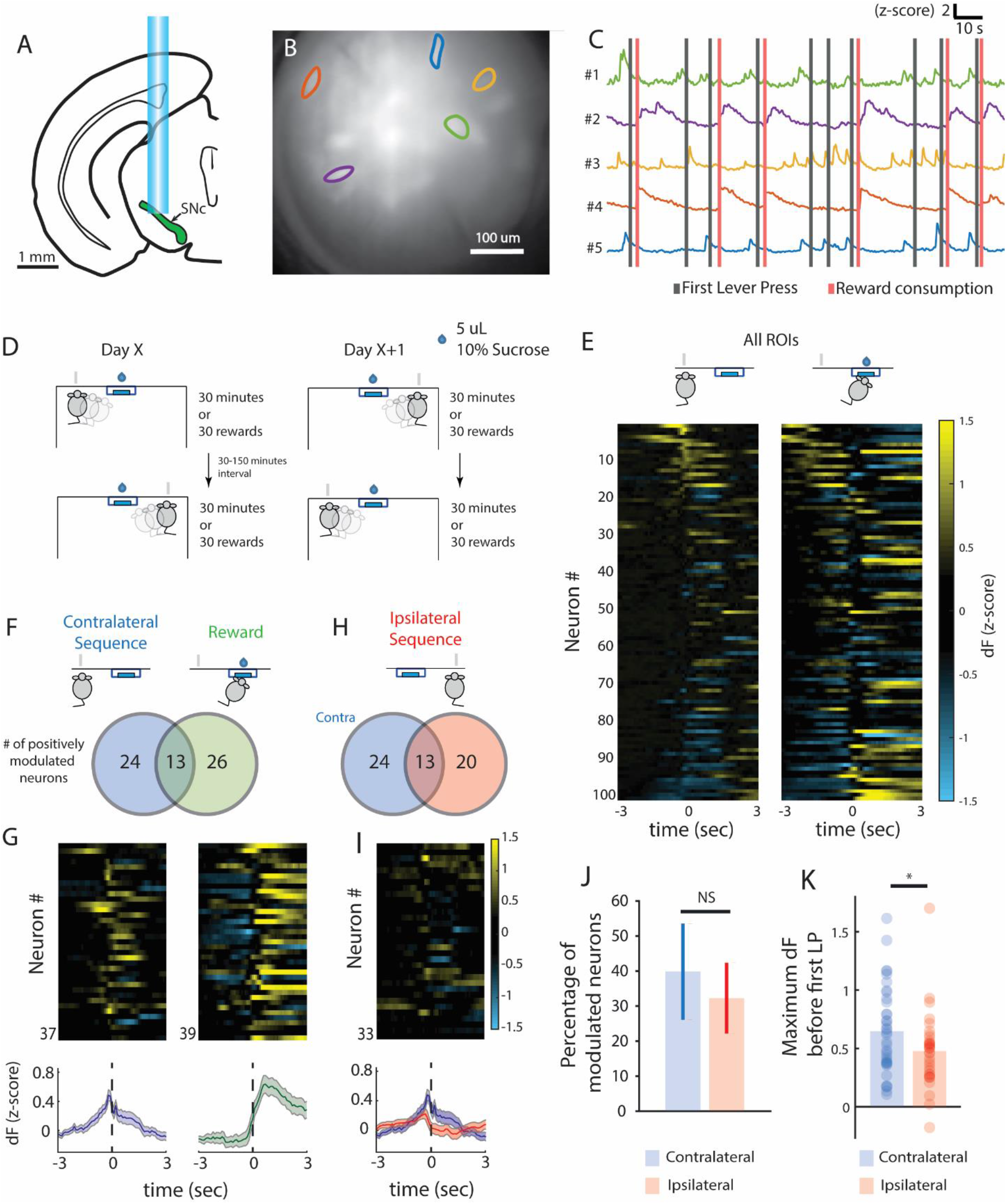
SNc dopaminergic neurons are transiently active before movement sequence initiation. **A)** In 6 DAT-IRES:Cre mice, a miniature epifluorescence microscope was used for deep-brain calcium imaging from SNc dopaminergic neurons. Imaging was performed after animals learned the task up to an asymptotic stage. **B)** Field of view (projection of pixel standard deviation) of a DAT-Cre mouse expressing GCaMP6f in the SNpc. Regions of interest (ROIs) correspond to traces in C. **C)** Example traces obtained using the CNMF-E algorithm during the FR4 task. ROIs #1 and #5 are examples of units modulated before first lever press - gray lines - and ROIs #2 and #4 are examples of units modulated after reward - red lines. **D)** Schematics of the training schedule used for this set of experiments. Mice were trained in a pseudo-randomized order across different training days alternating between starting with the ipsi- or the contralateral forelimb. **E)** Activity of all recorded ROIs from one session aligned to the first contralateral lever press (left) and to the beginning of reward consumption (right). **F)** Venn diagram representing the number of contralateral first press (left, blue color) and reward-related (right, green color) neurons. **G)** PETH of positively modulated neurons for first press and reward (bottom) and corresponding heat maps (top). **H)** Venn diagram representing first press modulated neurons when action was performed by contralateral (blue, same number as in F) and ipsilateral (red) forelimb. **I)** PETH of positively modulated neurons for first press contralateral (blue) and ipsilateral (red) (bottom) and corresponding heat map for ipsilateral neurons (top). **J)** Percentage of positively modulated neurons before first lever press per mouse, comparing contralateral and ipsilateral forelimb (NS, p = 0.736, paired t-test). **K)** Maximum fluorescence in the 2 seconds before first lever press of positively modulated neurons when the action was performed with the contralateral and ipsilateral forelimb (Contralateral, n=37 neurons, Ipsilateral, n=33 neurons, * p=0.0480, unpaired t-test). Data are presented as mean ± SEM. *p<0.05; For detailed statistical analysis, see Table S1.

Constrained non-negative matrix factorization for endoscope data (CNMF-E) (27) was used to extract activity traces for individual neurons from the microscope video (6 mice, 101 neurons; Fig 2B-C). Neuronal spatial footprints and temporal activity were extracted from conjoined left and right sessions. Then, for all subsequent analysis, a normalized version (z-score) of the scaled, non-denoised version of dF extracted by CNMF-E, was performed for each of the two full sessions (left and right) independently. We created peri-event time histograms using the normalized fluorescence for first press in any lever press sequence and reward consumption for each experimental condition (ipsilateral or contralateral forelimb performance Fig 2D-E).

In line with previous results (2), we found a that ~37% of SNc dopaminergic neurons were modulated before movement sequence initiation and a similar percentage was modulated around reward. The overlap between these 2 populations was small (~13%) and not different from what expected by chance (Fig 2F,G, Fig. S4). Some DANs also started their modulation during sequence execution, but this number was lower than those modulated before sequence performance (Only 7-13% of neurons modulated during performance, F(1,10)=17.33, p=0.009; Fig S5A).

Movement-initiation neurons displayed transient increase in activity before both contralateral and ipsilateral forelimb movement sequences (Fig 2H,I Fig S5B). However, although a similar proportion of movement-initiation neurons was active before contra and ipsilateral movements (39.6 +- 13.7% vs 32.1 +- 10.01% p=0.7356, paired t-test, Fig 2J), the magnitude of the activity of these neurons was significantly higher in the contralateral vs. ipsilateral SNc (0.646 +- 0.061 vs. 0.427 +- 0.059, p=0.048, unpaired t-test, Fig 2K). This difference in neural activity could not be explained by a difference in the performance of the task as no performance differences were identified between ipsi and contralateral forelimb (Fig. S3, Table S2).

These data show that although movement-initiation neurons were found bilaterally, activity preceding contralateral limb movements is higher that the activity preceding ipsilateral movements.

### DANs encode the vigor of upcoming contralateral, but not ipsilateral, forelimb movements

We next investigated the relation between the magnitude of the dopamine transients before movement initiation and the vigor of the sequences. The magnitude of activity of movement-modulated DANs was sorted according to the number of presses in each sequence for either ipsi and contralateral sequences. The maximum activity preceding sequence initiation was linearly related to the number of presses/sequence performed by the contralateral (Fig 3A left, n=37 neurons, p<0.001), but not ipsilateral forelimb (Fig 3A right, n=33 neurons, p=0.2660).

**Figure 3:**
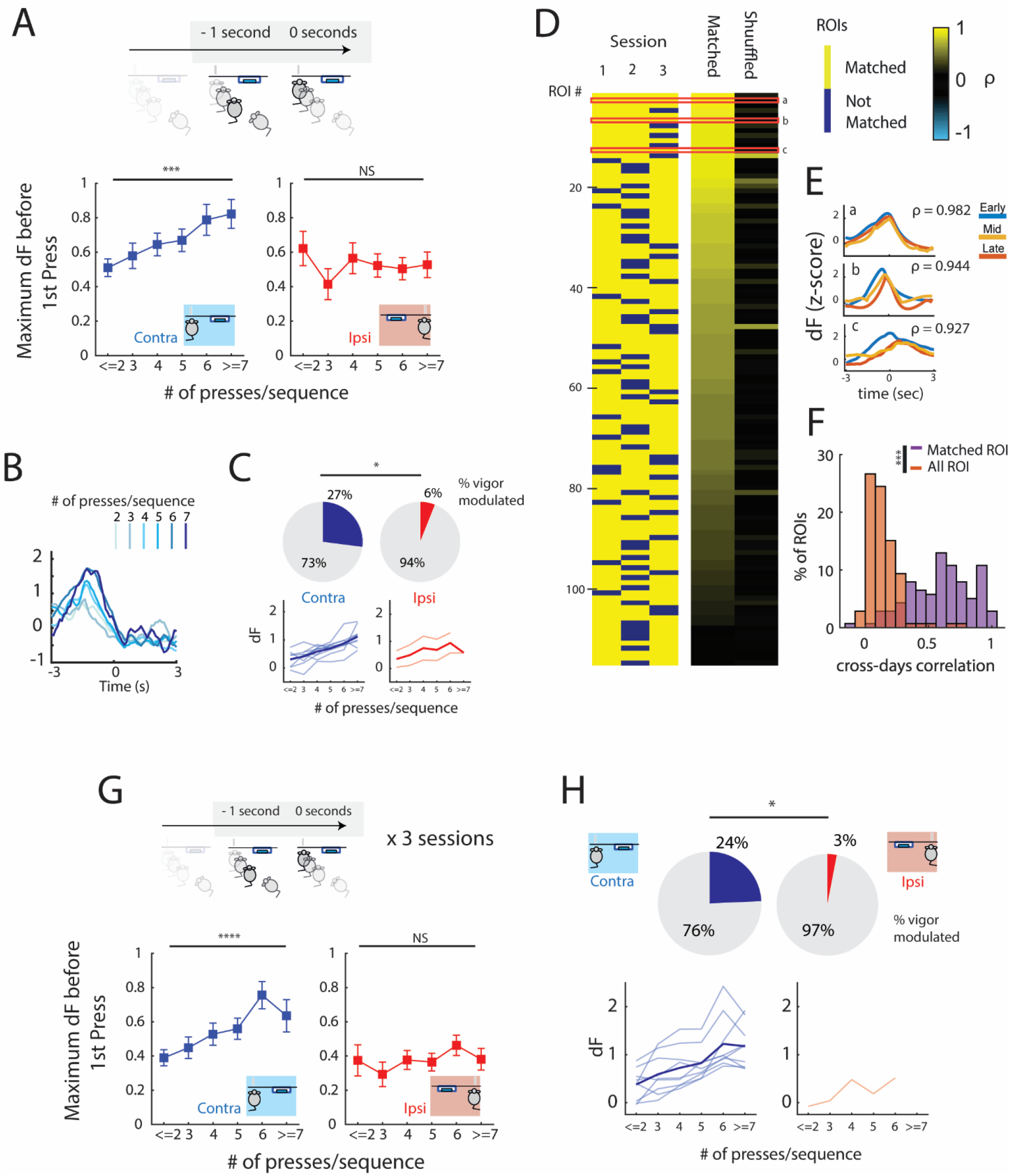
Transient SNc activity before the first lever press encodes the vigor of contralateral movement sequences. **A)** Maximum fluorescence before first lever press for movement-modulated neurons according to the number of presses/sequence performed by contralateral (left panel, n=37, F(1,168)=20.65, ***p<0.001, test for linear trend) and ipsilateral forelimb (right, n=33, F(1,151)=0.4521, p=0.5023, test for linear trend). **B)** Example of one movement-modulated neuron activity sorted by the number of presses/sequence. **C)** Percentage of neurons positively-modulated by vigor (top) in contra and ipsilateral conditions (*p=0.0266, Fisher’s exact test). Activity of individual neurons positively-modulated by vigor (light-color) and average activity (strong color) in contra and ipsilateral conditions (bottom). **D)** Matching of ROIs identified across 3 sessions of performance. Only ROIs that were matched in at least 2 of 3 sessions were plotted (left). Heatplot showing the average cross-days correlation of matched neurons PETHs and maximum cross-days correlation of each ROI with all ROIs of the same animal across days as a control (right). **E)** Example PETHs of 3 ROIs in different sessions disclosing the similarities in average neural activity. **F)** Histogram of the cross-days correlations represented in panel C). (n=114 ROIs *** p<0.001, paired t-test). **G)** Neurons were matched across 3 sessions. Maximum fluorescence 1 second before first lever press for movement-modulated neurons according to the number of presses/sequence performed by contralateral (left panel, n=37, F(1,175)=24.69, ****p<0.0001, test for linear trend) and ipsilateral forelimb (right, n=33, F(1,157)=1.202, p=0.2470, test for linear trend). **H)** Percentage of neurons positively-modulated by vigor (top) in contra and ipsilateral conditions across 3 days (*p=0.0151, Fisher’s exact test). Activity of individual neurons positively-modulated by vigor (light-color) and average activity (strong color) in contra and ipsilateral conditions (bottom).*p<0.05; **p<0.01; ***p<0.001; **** p<0.0001. Data are presented as mean ± SEM. For detailed statistical analysis, see Table S1

Having identified that specific neurons were modulated by vigor (Fig 3B, Fig S7E) we quantified how many were significantly modulated by vigor. We identified a significantly higher number of vigor-modulated neurons contralaterally (27%, Fig 3C left) than ipsilaterally (6%, Fig 3C right, p=0.0496). Negatively-modulated neurons were scarce (Fig S6A). This data supports the hypothesis that transient activity of a population of SNc DANs activity encode contralateral vigor.

Even accounting for some day-to-day variability in the field of view, *in vivo* calcium imaging permits us to track the same region of interest (ROI) across multiple sessions. This approach allows us to explore if SNc neurons’ activity is functionally stable across different performance sessions of highly-trained movements (supporting the argument of a functional identity of specific dopaminergic neurons during task performance in a context of diversity of functions – Fig 2F). To match neurons across sessions, we used a nearest neighbor approach. For all sessions, a centroid for each ROI was calculated. For each reference centroid, distance from all centroids on the image to be compared was calculated, and the 3 ROIs with the smallest distance were visually inspected for their shape to confirm the matching. Pairing was iteratively performed across the 3 included sessions. Across the 3 sessions, we identified 114 individual ROIs that were matched in at least 2 sessions (40 ROIs – 35.09% - were matched across the 3 sessions, Fig 3D)

Event-aligned activity of matched neurons was correlated across daily sessions and the results were averaged if matched more than 2 sessions (matched group). The activity of the same neuron was correlated with non-matched neurons from the same animal and averaged across daily sessions (shuffled group). This approach revealed that there is a high correlation of event-aligned activity of spatially mapped neurons (66.7% of neurons had a strong correlation – above 0.5, Fig 3D, Fig 3E) and this similarity was significantly higher than correlations with non-spatially mapped ones (0.677 +- 0.049 vs. 0.149 +- 0.014, paired t-test performed in the Fisher’s Z transformed correlation coefficient t=14.66, df=113, p<0.0001, Fig 3F). This analysis suggests that, in general, SNc dopaminergic neurons functional identity remains stable across days while animals perform the learned task.

Matching across days (as performed in Fig 3E) allowed us to study the activity of the same neuron during trials from different sessions (Fig S6B). For each neuron defined as movement-modulated we extracted the activity before movement initiation (as described in Fig 3A). Across the 3 sessions, a relationship with vigor was present contralaterally (Fig 3G, left, n=37 neurons, p<0.0001) but not ipsilaterally (Fig 3H, right, n=33 neurons, p=0.2470). A significant over-representation of vigor-modulated neurons contralaterally was observed across the 3 sessions (24% vs 3%, p=0.0181, Fig 3I)

These data show that activity in movement-initiation SNc neurons code the vigor of contralateral (in comparison to ipsilateral) performed movement sequences, as neural identity keeps stable over time.

### Reward-related dopamine activity is not lateralized

Modulation around reward was identifiable when mice performed the task either in the ipsi and contralateral conditions (Fig 4A,B). The number of neurons was not significantly different between conditions (Fig 4C, 36.9% +- 5.9 vs. 41.1 +- 14.1, paired t-test: p=0.8003, Fig 4D). The number of reward-modulated neurons (active only after reward consumption) was similar when the action leading to that reward was performed ipsi or contralaterally (~23%, Fig 4F middle, paired t-test: p=0.8902) and no difference was observed in the maximum fluorescence of neurons according to the side of performed action (0.95 +- 0.10 vs. 1.11 +- 0.11 Fig 4F right, t-test: p=0.3355). The overlap between reward-modulated neurons and movement initiation neurons was residual (~5% of the total) and significantly lower than the one we would expect by random allocation (Fig S7), revealing that movement initiation and reward neurons mostly represent two distinct populations.

**Figure 4:**
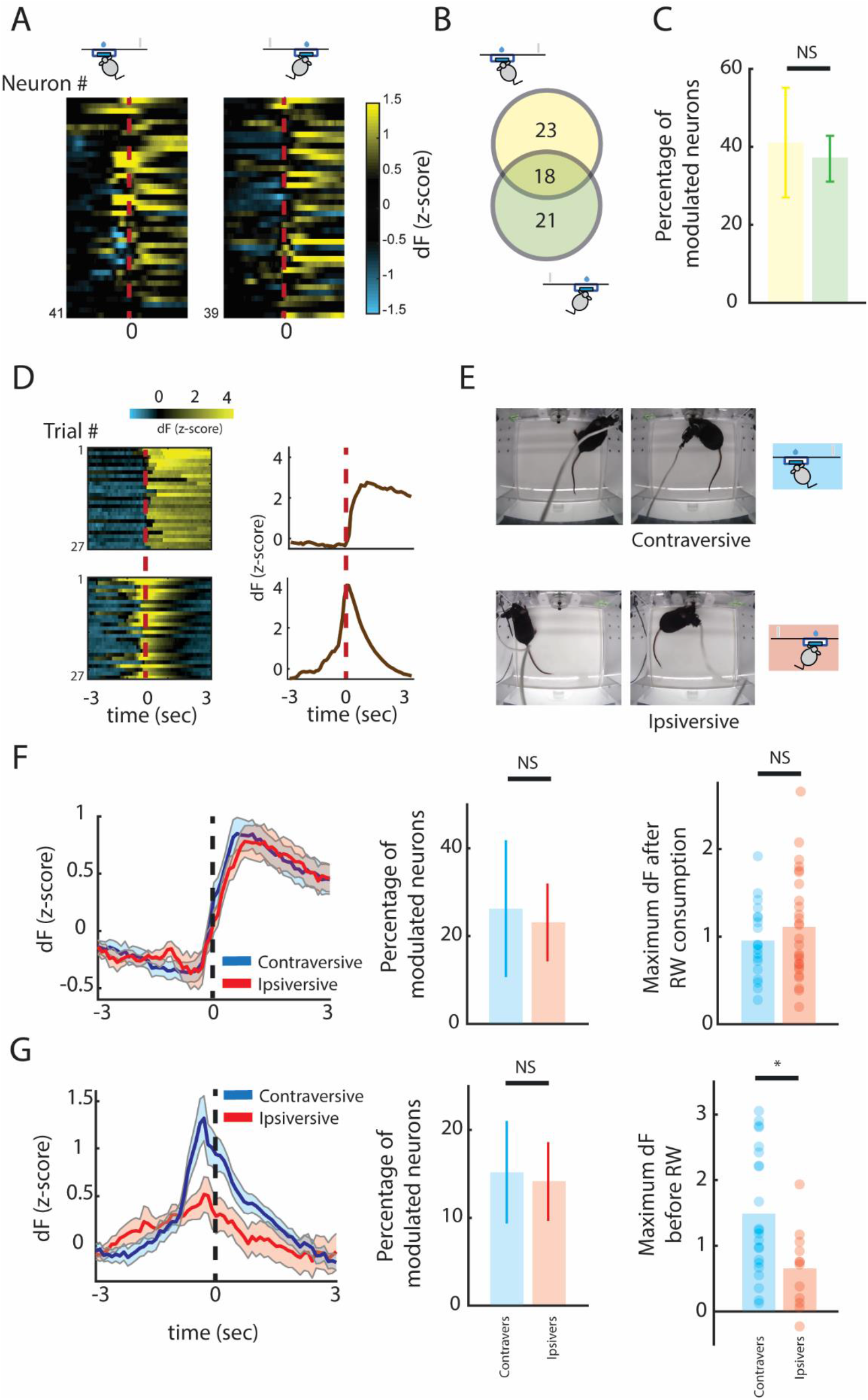
Activity of reward-modulated neurons is not lateralized. **A)** Heat maps of neurons with responses around reward when animals performed the task with the ipsilateral and contralateral forelimb. **B)** Venn diagram representing these neurons when reward is collected after performing a contraversive (yellow) and ipsiversive (blue, same number as in 2F) movement. **C)** Percentage of positively modulated neurons in the contra- and ipsiversive conditions (p = 0.80, paired t-test). Data are presented as mean ± SEM. **D)** Example of 2 neurons displaying different responses identified around reward: reward-modulated neuron (top) and magazine approach neuron (bottom). Left - Activity heatmap for all trials aligned to reward collection; Right - PETH of that neuron aligned to reward collection. **E)** Top view of mouse position when pressing the lever and consuming reward in situations where it performs a contraversive (top) and ipsiversive (bottom) movement. **F)** Activity of reward modulated neurons after a contraversive (blue) or ipsiversive (red) movement: PETH of reward modulated neurons (left), percentage of modulated neurons per mouse (n=6, 25.88% +- 15.40 vs. 22.80% +- 8.84, p=0.8902, paired t-test) and Maximum fluorescence after reward consumption (right) (Ipsiversive, n=27 neurons, Contraversive, n=19 neurons, p=0.3355, unpaired t-test). **G)** Activity of magazine-approach neurons after a contraversive (blue) or ipsiversive (red) movement: PETH of magazine-approach neurons (left), percentage of modulated neurons per mouse (n=6, 15.20% +- 5.85 vs. 14.14% +- 4.47, p=0.9125, paired t-test) and Maximum fluorescence (right) (Ipsiversive, n=12 neurons, Contraversive, n=22 neurons, *p=0.0104, unpaired t-test). *p<0.05. Data are presented as mean ± SEM. For detailed statistical analysis, see Table S1

A second group of neurons was already active before reward consumption and ramped up as animals approached the magazine (Fig 4D, bottom, hereafter called magazine-approach neurons) (28). Based on the box design, approach to the magazine could be performed with either an ipsi or a contraversive movement (Fig 4E). While the number of magazine-approach neurons was not significantly different during ipsi or contraversive movements (~15%, paired t-test: p=0.9125) their activity was significantly higher during contraversive movements (1.49 +- 0.21 vs. 0.66 +- 0.18 Fig 4G right, t-test: p=0.0104).

These data support that movement initiation and reward modulated neurons in the SNc are not likely the same DAN population and neurons responsive to reward consumption do not have a lateralized representation.

### Dopaminergic depletion reduces the vigor of contralaterally performed forelimb movement sequences

The results shown above suggest that SNc dopaminergic activity is asymmetric during single forelimb movements, and that its’ magnitude is related to the vigor of contra but not ipsilateral sequences. We therefore tested if unilateral loss of dopaminergic neurons in SNc would preferentially affect the vigor of contralateral sequences. To achieve that, we used unilateral striatal injection of 6-hydroxydopamine (6-OHDA), a neurotoxin that selectively affects dopaminergic neurons. It causes rapid degeneration of striatal terminals (within hours after treatment) and changes in SNc DANs markers and cell body structure and numbers are detectable already 3 days after lesion (29).

A new group of 14 mice was trained in the task as described in Fig 2D until they reached an asymptotic stage (FR4/1 second). After this, using a stereotaxic approach, injection of 2 uL of 6-OHDA (n=8) or saline (n=6) was performed in dorsolateral striatum in a randomly chosen side (left or right). Post-operative care was performed during the following 7 days and mice did not have access to the operant boxes. After this time period, mice were again placed in the operant boxes and had to perform the task to obtain reward using the same criteria as before treatment (FR4/1 second) (Fig 5A, B). Severity of dopamine depletion is revealed in Figs S9A-S9C.

**Figure 5:**
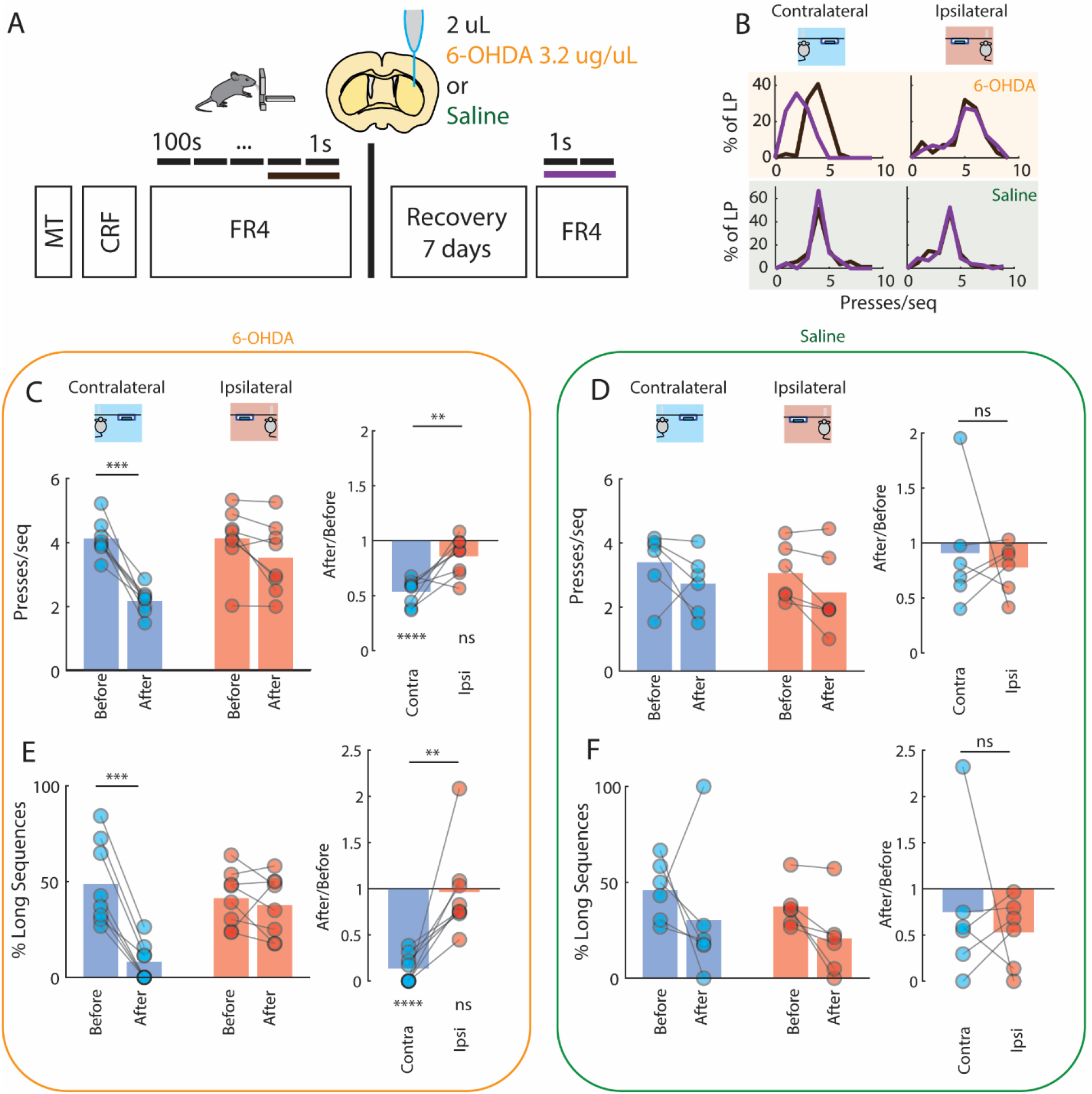
Dopamine depletion disrupts contralateral vigor. **A)** A new group of mice (n=14) was trained in the task performing actions with each forepaw. After a plateau performance was reached, mice were injected with 6-OHDA or saline unilaterally in the striatum and retested after the lesion. **B)** Example of performance with contra- and ipsilateral forelimbs to the lesion side, before (black) and after (purple) treatment of a mouse injected with 6-OHDA (top) and saline (bottom). Intra-striatal treatment with 6-OHDA leads to a redistribution of number of presses/sequence performed on the contralateral forelimb with the performance of sequences. **C)** Change in the number of presses/sequence for 6-OHDA treated animals. Left) Number of presses/sequence across the 4 conditions (Time: Before/After and Forelimb: Contra/Ipsilateral) for 6-OHDA treated animals. There was an effect for Time and an interaction between Forelimb and Time conditions. 2 Way repeated-measures ANOVA, Time F(1,7)=68.90, p<0.001, Forelimb F(1,7)=4.704, p=0.0667, Time x Forelimb F(1,7)=11.11, p=0.0125. Post-hoc tests revealed a significant difference in the before/after condition in the contralateral forelimb (4.12 +- 0.20 to 2.17 +-0,14, multiple comparison test after 2-way repeated-measures ANOVA, t(7)=6.835, p<0.001) but not ipsilaterally (4.13 +- 0.34 to 3.52 +- 0.39, multiple comparison test after 2-way repeated-measures ANOVA, t(7)=2.121, p=0.1380). Right) Ratio of presses/sequence after treatment, normalized to the one before treatment for both ipsi- and contralateral forelimbs (normalization of data presented on the left panel). There was a significant difference between ipsi and contralateral change (paired t-test, t(7)=3.759, p=0.0071) N=8. While contralaterally the change was significantly different from the unit value (one sample t-test vs. 1: t(7)=11.07, p<0.0001), ipsilaterally no significant difference was found (one sample t-test vs. 1: t(7)=2.281, p=0.0565). **D)** Change in the number of presses/sequence for saline treated animals. Left) Number of presses/sequence across the 4 conditions (Time: Before/After and Forelimb: Contra/Ipsilateral) for saline treated animals. There was only small effect for Time. 2 Way repeated-measures ANOVA, Time: F(1,5)=7.704, p=0.0391, Forelimb: F(1,5)=2.007, p=0.2157, Time x Forelimb: F(1,5)=0.01041, p=0.9227. There was not any significant change in the number of presses/sequence after saline treatment either contralaterally (3.40 +- 0.35 to 2.73 +- 0.33, multiple comparison test after 2-way repeated-measures ANOVA, t(5)=1.408, p=0.3885) or ipsilaterally (3.06 +- 0.31 to 2.46 +- 0.45, multiple comparison test after 2-way repeated-measures ANOVA, t(5)=1.264, p=0.4552). Right) Ratio of presses/sequence after treatment, normalized to the one before treatment for both ipsi- and contralateral forelimbs (normalization of data presented on the left panel). There wasn’t any significant difference between ipsi and contralateral change (paired t-test, t(5)=0.4441, p=0.6755). **E)** Change in the percentage of Long Sequences/All Sequences for 6-OHDA treated animals. Long Sequences are defined as sequences with a number of presses higher than the mean number of presses on baseline condition for each forelimb (i.e. the number calculated in panels C and D). Left) Percentage of Long Sequences/All Sequence across the 4 conditions (Time: Before/After and Forelimb: Contra/Ipsilateral) for 6-OHDA treated animals. There was an effect for Time and an interaction between Forelimb and Time conditions. 2 Way repeated-measures ANOVA, Time F(1,7)=30.12, p<0.001, Forelimb F(1,7)=2.087, p=0.1918, Time x Forelimb F(1,7)=32,45, p<0.001. Post-hoc tests revealed a significant difference in the before/after condition in the contralateral forelimb (48.82% +- 7.72 to 8.13% +- 3.48, t(7)=8.854, p<0.001) but not on ipsilateral forelimbs (41.34% +- 5.16 to 37.67% +- 5.67, t(7)=0.799, p=0.9725). Right) Ratio of long sequences after treatment, normalized to the one before treatment for ipsi- and contralateral forelimbs. There was a significant difference between ipsi and contralateral change (paired t-test, t(7)=4.126, p=0.0044). While contralaterally the change was significantly different from the unit value (one sample t-test vs. 1: t(7)=15.46, p<0.0001), ipsilaterally no significant difference was found (one sample t-test vs. 1: t(7)=0.210, p=0.8397). **F)** Change in the percentage of Long Sequences/All Sequences for saline treated animals. Left) Percentage of Long Sequences/All Sequence across the 4 conditions (Time: Before/After and Forelimb: Contra/Ipsilateral) for saline treated animals. There were no significant changes with saline treatment (2 Way repeated-measures ANOVA, Time F(1,5)=4.911, p=0.078, Side F(1,5)=0.8281, p=0.405, Time x Side F(1,5)=0.003, p=0.957; Contralateral: 45.87% +- 7.11 to 30.40% +- 14.40; Ipsilateral: 37.35% +- 4.72 to 20.69% +- 8.20). Right) Ratio of long sequences after treatment, normalized to the one before treatment for ipsi- and contralateral forelimbs. There wasn’t any significant difference between ipsi and contralateral change (paired t-test, t(5)=0.5099, p=0.6319). Data are presented as mean ± SEM. *p<0.05; **p<0.01; ***p<0.001; **** p<0.0001. For detailed statistical analysis, see Table S1.

Unilateral dopamine depletion led to a reduction in the number of presses per sequence (2 Way repeated-measures ANOVA, Time F(1,7)=68.90, p<0.001, Forelimb F(1,7)=4.704, p=0.0667, Time x Forelimb F(1,7)=11.11, p=0.0125, Fig 5C left) and corresponding reduction in the percentage of long/high vigor sequences (2 Way repeated-measures ANOVA, Time F(1,7)=30.12, p<0.001, Forelimb F(1,7)=2.087, p=0.1918, Time x Forelimb F(1,7)=32,45, p<0.001 Fig 5E left). Whereas the contralateral limb to the lesion started to perform smaller sequences (4.12 +- 0.20 to 2.17 +-0,14, Fig 5C left, Multiple comparison after 2-way repeated-measures ANOVA, p<0.001) and show a reduced number of long/high vigor sequences (48.82% +- 7.72 to 8.13% +- 3.48, Fig 5E left, Multiple comparison after 2-way repeated-measures ANOVA, p<0.001) this was not observed for the limb ipsilateral to the lesion (4.13 +- 0.34 to 3.52 +- 0.39, Fig 5C left, Multiple comparison after 2-way repeated-measures ANOVA, p=0.138 and 41.34% +- 5.16 to 37.67% +- 5.67, Fig 5E left, Multiple comparison after 2-way repeated-measures ANOVA, p=0.9725, Movies S3 and S4). Regardless of the unilateral dopamine depletion mice kept performing the task with the intended limb (Fig. S8).

The relative mean sequence length (after treatment/before treatment) was significantly different between sides (Fig 5C right, paired t-test, t=3,759, df=7 p=0.007) and significantly different from the unit value (representing no change) only contralaterally (Fig 5C right, one sample t-test, t=11.07, df=7, p<0.001). This result was confirmed by the significant difference in the relative proportion of long sequences between sides (Fig 5E right, paired t-test, t=4,126, df=7 p=0.004, significance from the unit value only identified contralaterally, Fig 5E right, one sample t-test, t=15.46, df=7, p<0.001)

By contrast, injection of saline only led to a small, and non-side specific reduction in the average number of presses/sequence (Fig 5D left, 2 Way repeated-measures ANOVA, Time F(1,5)=7.704, p=0.039, Side F(1,5)=2.007, p=0.216, Time x Side F(1,5)=0.010, p=0.923) without a significant change in the proportion of long sequences (Fig 5F left, 2 Way repeated-measures ANOVA, Time F(1,5)=4.911, p=0.078, Side F(1,5)=0.8281, p=0.405, Time x Side F(1,5)=0.003, p=0.957). When the relative mean sequence length was compared between sides no difference was found (Fig 5D right, paired t-test, t=0.44, df=5, p=0.6755), similar to the lack of a difference in sequence length change (Fig 5F right, paired t-test, t=0.5099, df=5 p=0.6319),

Overall, these data show that unilateral dopamine depletion leads to a reduction in the length of contralaterally performed movement sequences, without impacting the vigor or performance of ipsilateral movements.

## Discussion

We found that activity in a subset of SNc dopaminergic neurons encodes the vigor of contralateral actions before movement initiation. Using a novel lateralized task, we unraveled that transient SNc dopaminergic activity precedes the execution of forelimb movement sequences irrespective of the limb performing the sequence. However, this signal was only related to the duration of contralateral (but not ipsilateral) sequences. Also, striatal dopamine depletion disrupted the duration of contralateral, but not ipsilateral, movement sequences. By contrast, responses of reward-related SNc neurons were bilateral and modulated irrespectively of the side of the preceding action.

Laterality is a major topic in nervous system organization, and most attention on nigro-striatal pathway has been placed on the dopaminergic striatal terminals. Although in freely moving animals, striatal dopamine transients are synchronized across hemispheres (30), contralateral action response was identified in DA terminals in dorsal striatum (7). SNc neurons have both ipsilateral and contralateral functionally relevant projections (31), but those ipsilateral to SNc cell bodies are anatomically overrepresented (32). Our data suggest that activity in DANs cell bodies is lateralized, and hence the higher activity in striatal DA terminals observed before contralateral movements are not solely explained by more arborization of SNc neurons to ipsilateral striatum, or enhanced terminal modulation of DA release. Furthermore, and given that dopamine depletion starts asymmetrically by the caudal putamen in PD (19, 33), these findings have implications for understanding the asymmetry in movement vigor observed in PD (12, 34).

It has been previously shown that dopamine and its’ metabolite 3,4-dihydroxyphenylacetic acid (DOPAC) increase bilaterally in the striatum in relation to speed when rodents ran straight, but contralaterally when they performed circling movements (35). This is in line with the asymmetry in SNc neuronal activation we observed, and suggests that asymmetric tonic levels of dopamine in dorsal striatum regulate movement vigor (36). Furthermore, our results suggest that transient changes in SNc dopamine preceding movement onset, in addition to tonic activity, modulate kinematic aspects of contralateral movements.

Although evidence of laterality is present in movement responses, the same is not true for reward responses. This activity was similar irrespective of the side of the performed action that led to it and started only after reward collection. The lack of laterality of reward responses was also noted in a fiber photometry study in ventral striatum where headfixed mice had to perform an ipsi or contralateral movement in response to a visual stimulus (37). This suggests that responses to unpredicted reward represent a more general teaching signal in the brain, and not solely related to the action performed to obtain the reward.

Although efforts have been made to unify observations of movement and reward responses in DANs, under classic RPE model, some studies suggest that movement signals are modulated distinctly from RPE (38). The results presented here also suggest that movement signals and reward-related signals can be independently modulated. If movement-related signals would reflect learned action value, we would not expect a difference in magnitude of activity based on which limb was used to perform the actions - as reward prediction is similar for both left and right limb performance of action sequences. Furthermore, we would not expect that magazine-approach activity (28) would be higher for contraversive than ipsiversive movements, as again these actions have the same expected value. These observations do not argue that dopamine neurons do not encode an RPE, just that not all dopamine neurons necessarily encode an RPE. Facing the high dimensionality of an organism behavior, error signals could be differently decomposed according to the condition (39) or different state information may be carried by dissociable parallel circuits (40, 41). However, they substantiate the existence of a population of SNc DA neurons directly involved in movement and movement invigoration.

The findings here deserve further expansion regarding some aspect of PD etiopathogenesis. First, the involvement of distinct SNc populations in movement initiation vs. reward - an observation previously described at axonal terminals in the striatum, (1) - is consistent with the notion of selective vulnerability of nigrostriatal degeneration and the origin of motor versus neuropsychiatric manifestations such as depression, anxiety or apathy (42). Second, the finding that SNc activity is directly linked with the execution and vigor of a learned movement sequence implicates that, contrary to classic understanding (43), the nigrostriatal dopaminergic system may continue to be activated and engaged during the performance of routine, automatic actions. Such neurons in the ventrolateral SNc are the first and most affected in PD and accordingly, the findings here agree with the hypothesis of high metabolic demand and overuse as critical vulnerability factor underlying the onset of neurodegeneration (44). Thus, clarifying if there is a link between genetic heterogeneity and functional phenotypes in these dopaminergic neurons could provide valuable resources to better understand the spectrum of clinical manifestations in PD and, more importantly, to define the origin of selective neuronal dopaminergic degeneration in PD.

## Materials and Methods

### Experimental Model and Subject Details

All experiments were approved by the Portuguese Direcção Geral de Veterinária and Champalimaud Centre for the Unknown Ethical Committee and performed in accordance with European Union Directive for Protection of Vertebrates Used for Experimental and other Scientific Ends (86/609/CEE and Law No. 0421/000/000/2014). Male C57BL/6J mice were tested between 2 and 4 months old. For calcium imaging studies, the male DAT-IRES:Cre (Dopamine Transporter-Internal Ribosome Entry Site-linked Cre recombinase gene) mouse line from Jackson Labs Stock 006660 (The Jackson Laboratory; B6.SJL-Slc6a3tm1.1(Cre)Bkmn/J) was used. These mice have Cre recombinase expression directed to dopaminergic neurons, without disrupting endogenous dopamine transporter expression. These studies were only performed in male mice, thus limiting generalization to female animals. Genotype was confirmed by polymerase chain reaction (PCR) amplification. Sample sizes are detailed in the Results and/or figure legends.

### Virus injections and lens placement

Mice were kept in deep anaesthesia using a mixture of isoflurane and oxygen (1-3% isoflurane at 1l/min) and the procedure was conducted in aseptic conditions.

The mouse head was stabilized in the stereotaxic apparatus (Koft), a skin incision was performed to expose the skull, connective and muscle tissue was carefully removed and the skull surface was leveled at less than 0.05mm by comparing the height of bregma and lambda, and also in medial-lateral directions. Unilateral virus injection was performed using a glass pipette with GCaMP6f stock viral solution (AAV2/5.SYN.FlexGCaMP6fWPRE.SV40 - University of Pennsylvania). For imaging, 1 ul of virus solution was injected in the right (n=3) of the left (n=3) substantia nigra compacta at the following coordinates: −3.16 mm anteroposterior, 1.40mm lateral from bregma and 4.20 deep from the brain surface. The injection was done using a Nanojet II or Nanojet III (Drummond Scientific) with a rate of injection of 4.6 nl every 5s. After the injection was finished, the pipette was left in place for 10-15 minutes. The virus solution was kept at −80 °C and thawed at room temperature just before the injection.

A 500-um diameter, 8.2-mm long gradient index (GRIN) lens (GLP-0584, Inscopix) was implanted at the same coordinates as the injection. Before the lens was lowered, a blunt 28 G needle was lowered to 3 mm deep from the brain surface to facilitate the lowering of the GRIN lens. The GRIN lens was then lowered (4.2 mm deep). The lens was fixed in place using cyanoacrilate, quick adhesive cement (C&B Metabond) and black dental cement (Ortho-Jet). Three weeks after surgery, the mouse was anaesthetized and fixed with head bars. A baseplante (BPC-2, Inscopix) attached to a mini epifluorescence microscope (nVista HD, Inscopix) was positioned above the GRIN lens. To correctly position the baseplate, brain tissue was imaged through the lens to find the appropriate focal plane using 40% LED power, a frame rate of 10 Hz and a digital gain of 4. Once the focal plane was set, the baseplate was cemented to the rest of the cap using the same dental cement. Imaging started 2–3 days after this final step.

### Single-limb fast FR4 operant task

Animals were trained using 14×16 cm custom-built operant chambers placed inside sound attenuating boxes. PyControl, (https://pycontrol.readthedocs.io), a behavioral experiment control system built around the Micropython microcontroller, was used to control and detect events and supply rewards. The custom-built boxes had in their design a retractable lever.

At the beginning of each session there was the onset of a light, and the animals were required to perform a sequence of presses at a minimum frequency in order to obtain a sucrose reward. Sucrose solution (10%) was delivered through the opening of a solenoid (LHDA1231515H, Lee Company). Sucrose solution was delivered through a tube into the magazine (5μl per reward). Licks were detected using an infrared beam and through a side camera, mouse position in the box was monitored through a camera placed on the top of the box.

Mice were placed on food restriction throughout training, and fed daily after the training sessions with approximately 1.5 - 2.5g of regular food to allow them to maintain a body weight of around 85% of their baseline weight. To facilitate learning, animals were initially exposed to one session of magazine training where sucrose would be available on a random time schedule, and to three to four sessions of continuous reinforcement schedule (CRF) before training, where single lever presses would be reinforced. In the following sessions animals were reinforced if they performed a sequence of 4 consecutive presses (Fixed Ratio 4, FR4) in a particular time window (FR4/Xs, fixed-ration four within X seconds). The duration of time required to perform the four lever presses was reduced across sessions from 100 seconds to 20 s, 8 s, 4 s, 2 s and finally 1s. To shape animals to use only one of the forelimbs the lever was progressively retracted and the slit thought which the forelimb accessed the lever was reduced with a custom-built piece.

In the imaging group, animals performed the task 2 times/session - one with the lever in the left side of the box and the other with the lever in the right - that were randomized throughout the training. Task ended after 30 minutes on each side or when the animals obtained 30 rewards.

The lever was equipped with a digital 9-axis inertial sensor with a sampling rate of 200 Hz (MPU-9150, Invensense) assembled on a custom-made PCB and connected to a computer via a custom-made USB interface PCB (Champalimaud Foundation Hardware Platform). Lever velocity was extracted from this sensor.

Timestamps from the behavioral task were synchronized with calcium imaging data using TTL pulses sent from the behavioral chambers to the Inscopix data acquisition system via a BNC cable.

### GCaMP6f imaging using a mini-epifluorescence microscope

Mice were briefly anaesthetized using a mixture of isoflurane and oxygen (1% isoflurane at 1L/min) and the mini-epifluorescence microscope was attached to the baseplate. This was followed by a period of 15-20 min of recovery in the home cage before starting the experiments. Fluorescence images were acquired at 10 Hz and the LED power was set 40-60% with a gain of 4. Image acquisition parameters were always set to the same parameters between sessions to be able to compare the activity recorded. Six GCaMP6f-expressing DAT-Cre mice were imaged during the FR4/1s task in 3 performance days.

### Calcium image processing and analysis

#### GCaMP6f image processing

All fluorescence movies were initially processed using the Mosaic Software (v. 1.2.0, Inscopix). Two different movies were collected on the same day (one for ipsi and one for contralateral forelimb). As the epifluorescence microscope was not removed during this period, movies were concatenated for the next analysis step. First, all frames were spatially binned by a factor of 4. To correct the movie for translational movements and rotations, frames were registered to a reference image consisting of an average of the raw fluorescence movie.

#### Extraction of calcium signals

We implemented the ‘constrained non-negative matrix factorization for endoscopic data’ (CNMF-E) framework for our calcium imaging analysis. This framework is an adaptation of the CNMF algorithm that can reliably deal with the large fluctuating background from multiple sources in the data, and enable accurate source extraction of cellular signals. It include four steps: 1) initialize spatial and temporal components of single neurons without the direct estimation of the background, 2) estimate the background given the estimated spatiotemporal activity of the neurons; 3) update the spatial and temporal components of all neurons while fixing the estimated background fluctuation, 4) iteratively repeat step 2 and 3.

CNMF-E only identifies regions of interest that are active in the condition.

After analysis, data from the videos were separated in ipsi and contralateral videos. Further calcium imaging analyses were performed on standardized scores (z-score) of each session.

#### Criteria to identify lever-press-related and reward-related DANs using GCaMP6f imaging

We constructed a PETH for each neuron trace spanning from −8 to 6 s from lever press onset for the first press and for the first lick after reward. Distributions of the PETH from −8 to −3 s before the event were considered baseline activity. We then searched each PETH during a determined epoch for bins that were significantly different from the baseline. A significant change in fluorescence was defined as at least two consecutive bins with fluorescence higher than a threshold of 99% above the baseline. For first-press modulated neurons, a window from −2 to 0 s was used. For rewarded lick modulated neurons a window from 0 to 1 s was used. For each neuron, maximum activity in specific time-windows was calculated by the maximum of a moving average of 3 bins.

#### Cell pairing across sessions

Analysis of matched cells between different days/sessions was based on a nearest neighbors method. For all sessions, a centroid for each ROI was calculated. For each reference centroid, distance from all centroids on the image to be compared was calculated, and the 3 ROIs with the smallest distance were visually inspected for their shape to define a match. Alignments were performed to 4 events: First lever press and Reward in the ipsi and contralateral situations. The calcium-signal from −10 to +6 seconds after the event was used to calculate the correlation coefficients for individual ROIs across days in the 4 conditions. If the neuron was matched in the 3 days, the 3 possible correlation coefficients were calculated and averaged. Then the maximum value was extracted. As a control, we correlated each ROI with all ROIs in the field of view (FOV) of the same animal on the comparison day. The maximum value was extracted and then averaged across units. Correlation values were transformed for each point using Fisher’s Z statistic, then the samples were averaged, and back-transformed into a weighted correlation.

#### PETH correlation across sessions

For each ROI, 4 PETHs were built as previously described: For the ipsilateral and contralateral conditions 2 events were considered (lever press and rewarded licks). For each ROI pair the Spearman correlation coefficient between these 4 events was calculated. The one with the maximum correlation was identified and this value extracted. This process was repeated when matching occurred across the 3 sessions, and average value between the 3 matchings (A and B, A and C and B and C) was computed. These are the values for matched ROIs in figure 3D.

As a control, we ran the same analysis, but instead of using the matched ROI we used all ROIs from the same animal, in a different session. We then calculated the average of the maximum correlations as previously described.

#### Sequence length analysis

A movement sequence was defined as a bout of lever presses occurring while the animal’s snout in spatially defined region of interest near the lever. For imaging experiments, as the cable could cause some distortion on head position estimate, we confirmed these results by defining a second ROI using the side camera. Sequences were divided by the number of presses performed. We grouped together sequences composed of 1 or 2 presses as sequences with 7 or more presses.

A neuron was classified as vigor related if the non-parametric correlation between its’ maximum activity and number of presses was significant (p<0.05).

### Striatal injection of 6-OHDA or Saline and post-operative care

As in virus injections, mice were kept in deep anaesthesia using a mixture of isoflurane and oxygen (1-3% isoflurane at 1l/min) and the procedure was conducted in aseptic conditions.

The mouse head was stabilized in the stereotaxic apparatus (Koft), a skin incision was performed to expose the skull, connective and muscle tissue were carefully removed and the skull surface was leveled at less than 0.05mm by comparing the height of bregma and lambda, and also in medial-lateral directions. 6-Hydroxydopamine hydrochloride (Sigma Aldrich AB, Sweden) was dissolved at a fixed concentration of 3.2 μg/μl free-base in 0.02% ice-cold ascorbic acid/saline and used within 2 h. Injection of 2 ul of 6-OHDA or saline was performed in the dorsolateral striatum +0.5 mm anteroposterior and ±2.5 mm lateral from bregma and 3.0 mm deep from brain surface. Injection was done through a glass pipette using a Nanojet II with a rate of injection of 4.6 nl every 5s. After the injection was finished, the pipette was left in place for 10-15 minutes.

After surgery, food restriction was reduced (with each animal having access to up 4 mg of food pellets/day). Mice that showed weight loss were hand-fed (i.e. they were presented with the food while being held by the hands of the investigator) and DietGel Boost was placed in their boxed up to 4 days after surgery. In order to avoid competition for the food, weaker mice were placed in cages other than those containing unimpaired mice. The postoperative survival rate was 100%.

#### Sequence length analysis

Movement sequences were divided into short and long sequences according to mean number of presses/sequence before striatal injection of 6-OHDA or Saline.

### Anatomical verification

Animals were euthanized after completion of the behavioural tests. First animals were anaesthetized with isoflurane, followed by intraperitoneal injection of ketamine-xylazine (5 mg/kg xylazine; 100 mg/kg ketamine). Animals were then perfused with 1% phosphate buffered saline (PBS) and 4% paraformaldehyde and brains were extracted for histological processing. Brains were kept in 4% paraformaldehyde overnight and then transferred to 1x PBS solution. Brains were sectioned coronally in 50-um slices (using a Leica vibratome VT1000S) and kept in PBS before mounting or immunostaining.

Images were taken using a wide-field fluorescence microscope (Zeiss AxioImager).

### Statistical analysis

Data is presented as mean ± standard error of mean (SEM) and statistical significance was considered for p < 0.05. Statistical analysis was conducted using GraphPad Prism 8 (GraphPad Software Inc., CA) and MATLAB statistical toolbox (The MathWorks Inc, MA). One-way or two-way ANOVAs were used to investigate main effects, and Šídák-corrected post-hoc comparisons performed whenever appropriate. Paired or unpaired t-tests were used for planned comparisons. For Figure 3, Spearman’s rank correlations were calculated. Details for statistical tests are presented in Table S1. Statistical methods were not used to pre-determine sample size

## Acknowledgments

We thank Ana Vaz and Catarina Carvalho for mouse colony management, Thomas Akam and Hélio Rodrigues for help in behavioral box development and implementation, and the Champalimaud Hardware Platform (Filipe Carvalho, Artur Silva and Dário Bento) for support in the development of the behavioral hardware setup. We thank Cristina Alcácer and Nuno Loureiro for their contributions during the 6-OHDA experiments. Further support was obtained from the research infrastructure Congento, co-funded by Lisboa2020 and FCT (LISBOA-01-0145-FEDER-022170).

## Funding

Fundação para a Ciência e Tecnologia (FCT) doctoral fellowship SFRH/BD/119623/2016 (MDM)

Fundação Luso-Americana para o Desenvolvimento (FLAD) visiting student fellowship 2018/31 (MDM)

Gulbenkian Foundation doctoral fellowship (JAdS),

Marie Curie Fellowship MSCA-IF-RI 2016 (LFH)

Spanish Ministry of Innovation and Science BES-2016-077493 (IC)

ERA-NET (RMC)

European Research Council COG 617142 (RMC)

HHMI IEC 55007415 (RMC)

National Institute of Health 5U19NS104649 (RMC)

Simons-Emory International Consortium on Motor Control (RMC).

## Author contributions

Conceptualization: MDM, JAdS, RMC

Investigation: MDM, JAdS, LH, IC

Formal analysis: MDM

Visualization: MDM

Supervision: RMC, JO

Writing—original draft: MDM, RMC

Writing—review & editing: MDM, RMC, JAdS, LH, IC, JO

## Competing interests

Authors declare that they have no competing interests

## Data and materials availability

All data needed to evaluate the conclusions in the paper are present in the paper and/or the Supplementary Materials. Data that support the findings of this report are available from the corresponding author upon reasonable request.

## Notes

### Competing Interest Statement

The authors have declared no competing interest.

