## Supplementary for "Transient dopamine neuron activity precedes and encodes the vigor of contralateral movements"

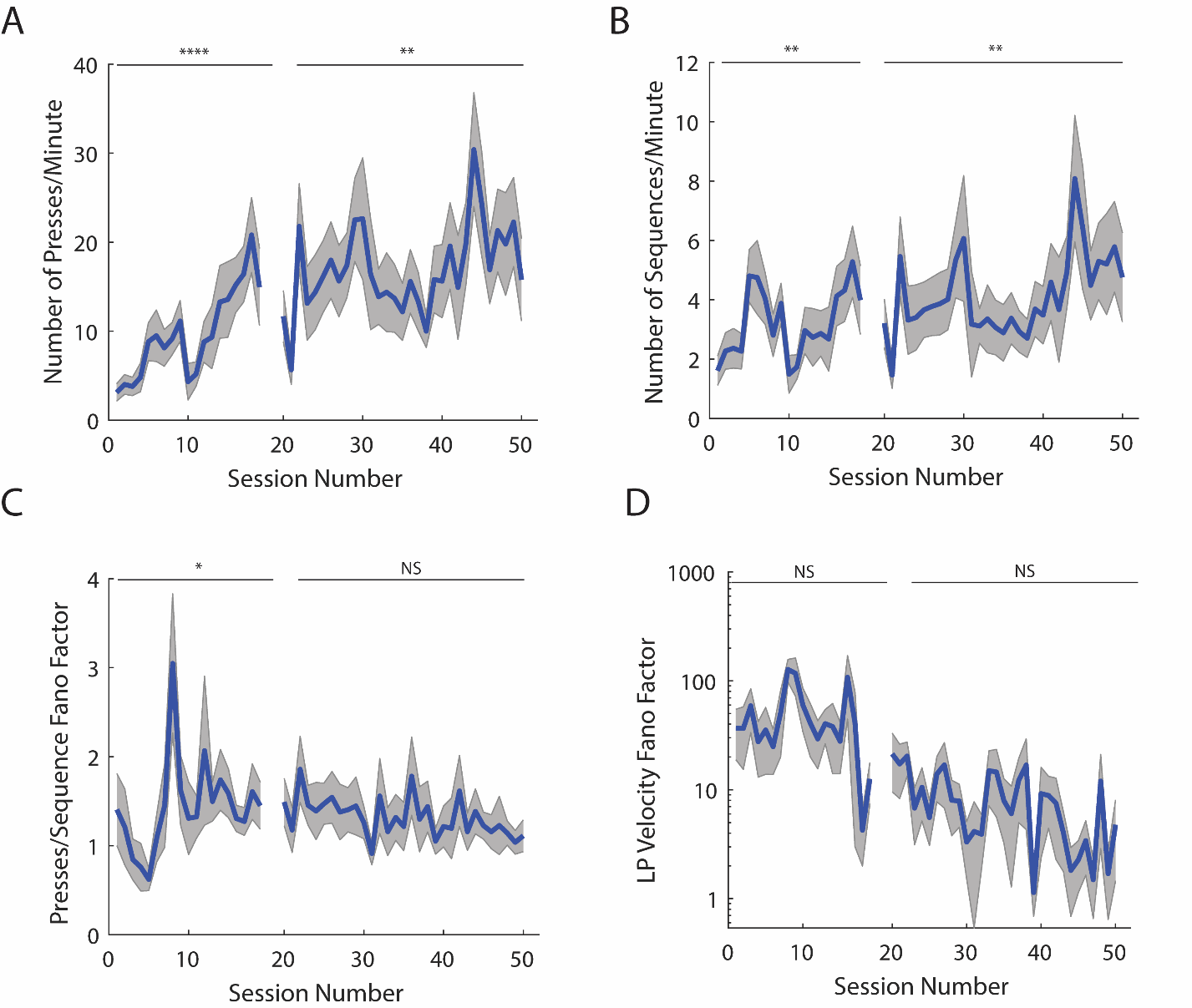

Fig. S1.

**A novel task for assessment of single individual forelimb movements.** **A)** Number of lever presses/minute. **B)** Number of performed sequences/minute. **C)** Fano Factor of the number of presses/sequence **D)** Fano Factor of the average velocity of lever press. Details on Table S2.

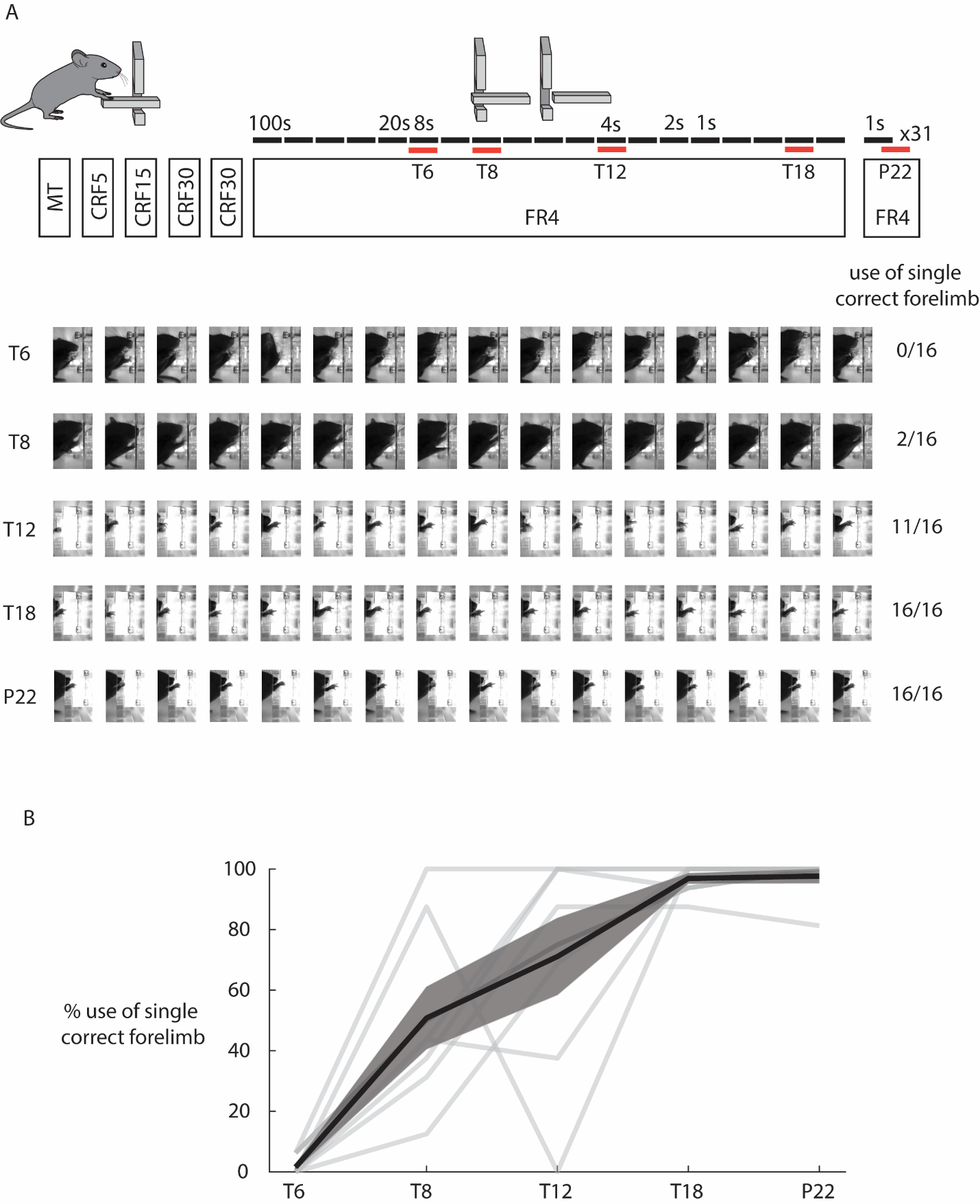

Fig. S2.

**A novel task for assessment of single individual forelimb movements. A)** Frames corresponding to the moment of lever press were identified for 5 specific session of the training schedule (Training days – T6, T8, T12 and T18 and Performance session – P22). Sixteen (16) randomly selected frames corresponding to 16 lever presses (or all, if less) were subsampled and visually inspected to assess the use of the correct forelimb. Data from one example animal is provided. **B)** Data from the 8 mice is summarized. In the last sessions near all presses (97.66% +- 2.34) were performed by the correct forelimb.

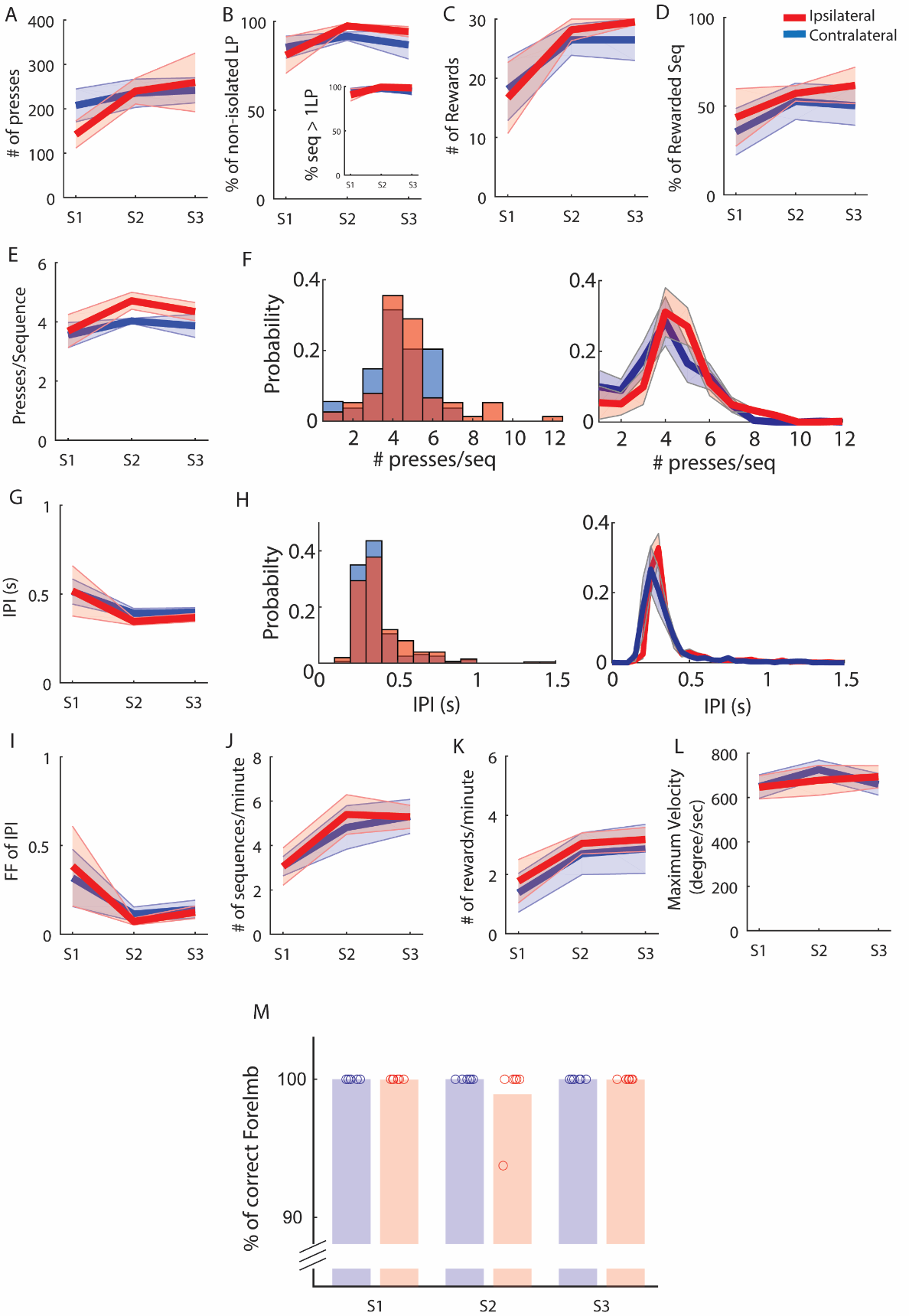

Fig. S3.

**Mice perform the task using either the ipsi and contralateral forelimbs.** Behavioral results from the 6 mice used in the analysis described in Figures 2-4. Data from sessions included in analysis in Figure 2 and Figure 3. Statistical details on Supplementary Table 2. **A)** Total number of presses/session, **B)** % of sequences composed by more than one lever press - Inset % non isolated lever presses **C)** Number of Presses/Sequence. **D)** Histogram of the distribution of number of presses/sequence for ipsi and contralateral movements for one example animal (left) and for all animals across one day (right). **E)** Number of Rewards/Session **F)** % of rewarded Sequences. **G)** InterPress Intervals. **H)** Histogram of the distribution of IPIs for ipsi and contralateral movements for one example animal (left) and for all animals across one day (right) **I)** Fano Factor of InterPress Intervals **J)** Number of sequences performed/minute **K)** Number of rewards obtained/minute. **L)** Maximum velocity per press. **M)** % of correct forelimb use.

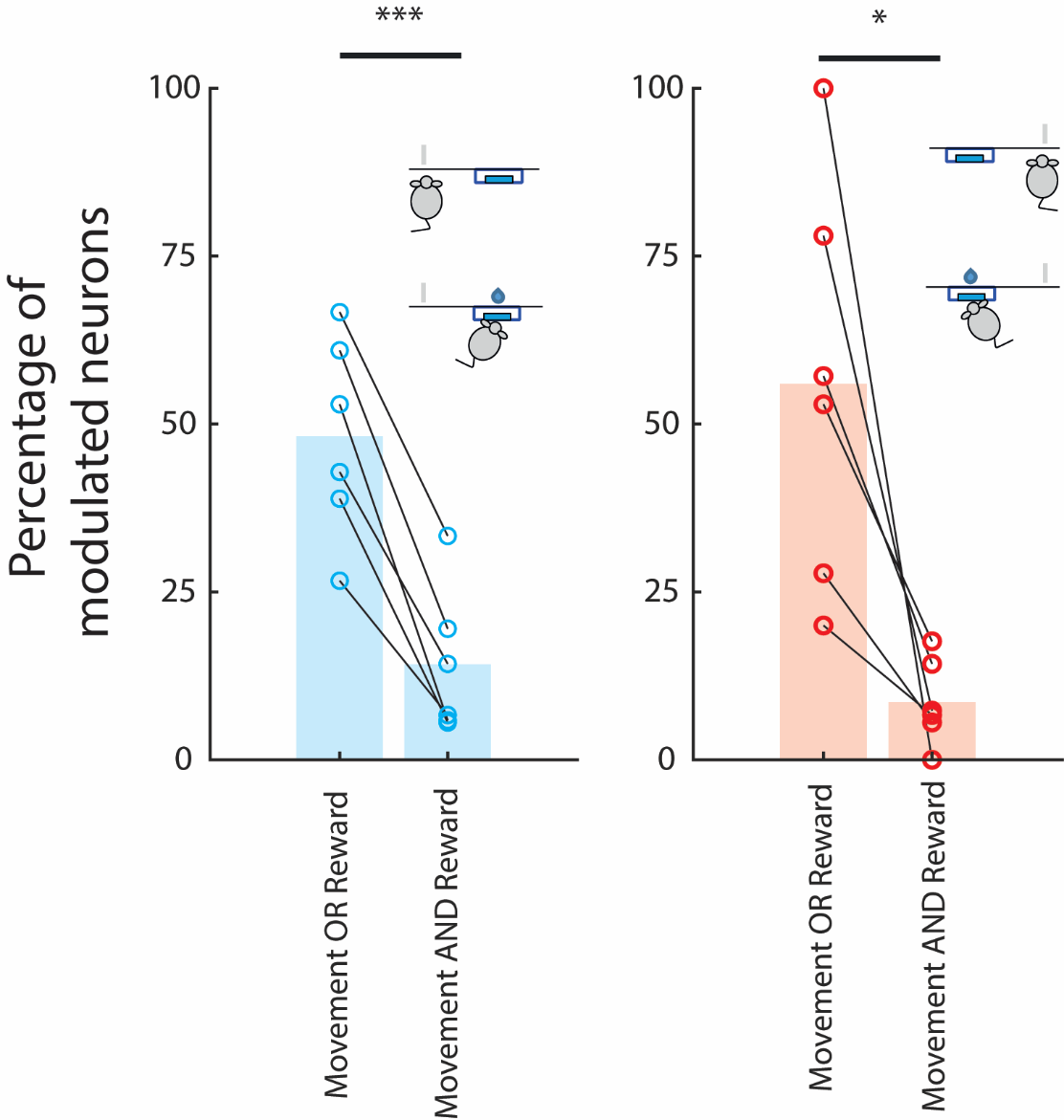

Fig. S4.

**SNc DANs are modulated by one event.** Neurons are more commonly modulated by only one event (movement or reward) than the two events (Left: 48,17 +- 6.07 vs. 14.21 +- 4.45, t=8.743, df=5, p<0.001; Right: 55.99 +- 12,28 vs 8.58 +- 2.60, t=3.576, df=5, p=0.0159)

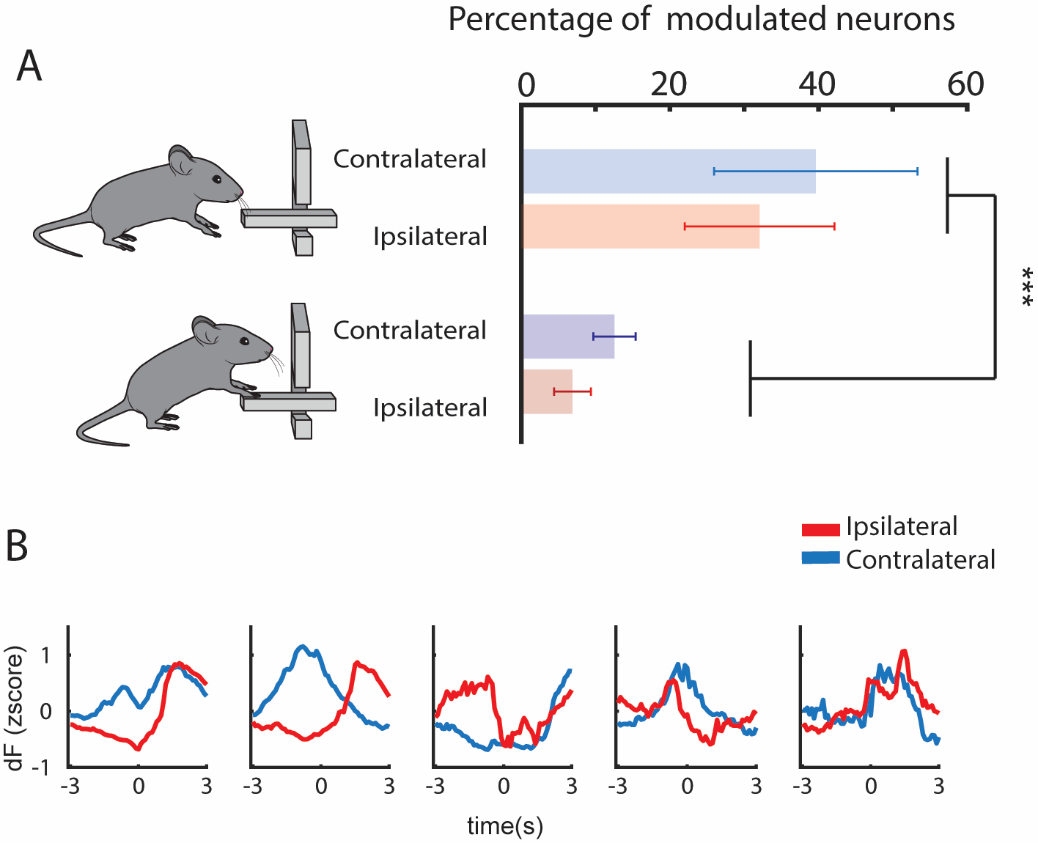

**Fig. S5.**

**Activity before movement onset is more common that during execution** **A)** Number of neurons whose modulation started before movement sequence initiation (compare with Figure 2) and during sequence execution. Two-way repeated-measures ANOVA; main effect before/after F(1,5)=17.33, p=0.0088; main effect side F(1,5)=0.3986, p=0.5555; interaction effect F(1,5)=0.008, p=0.9336 **B)** Example of matched ROIs aligned to first lever press when action is performed by contralateral and ipsilateral forelimb

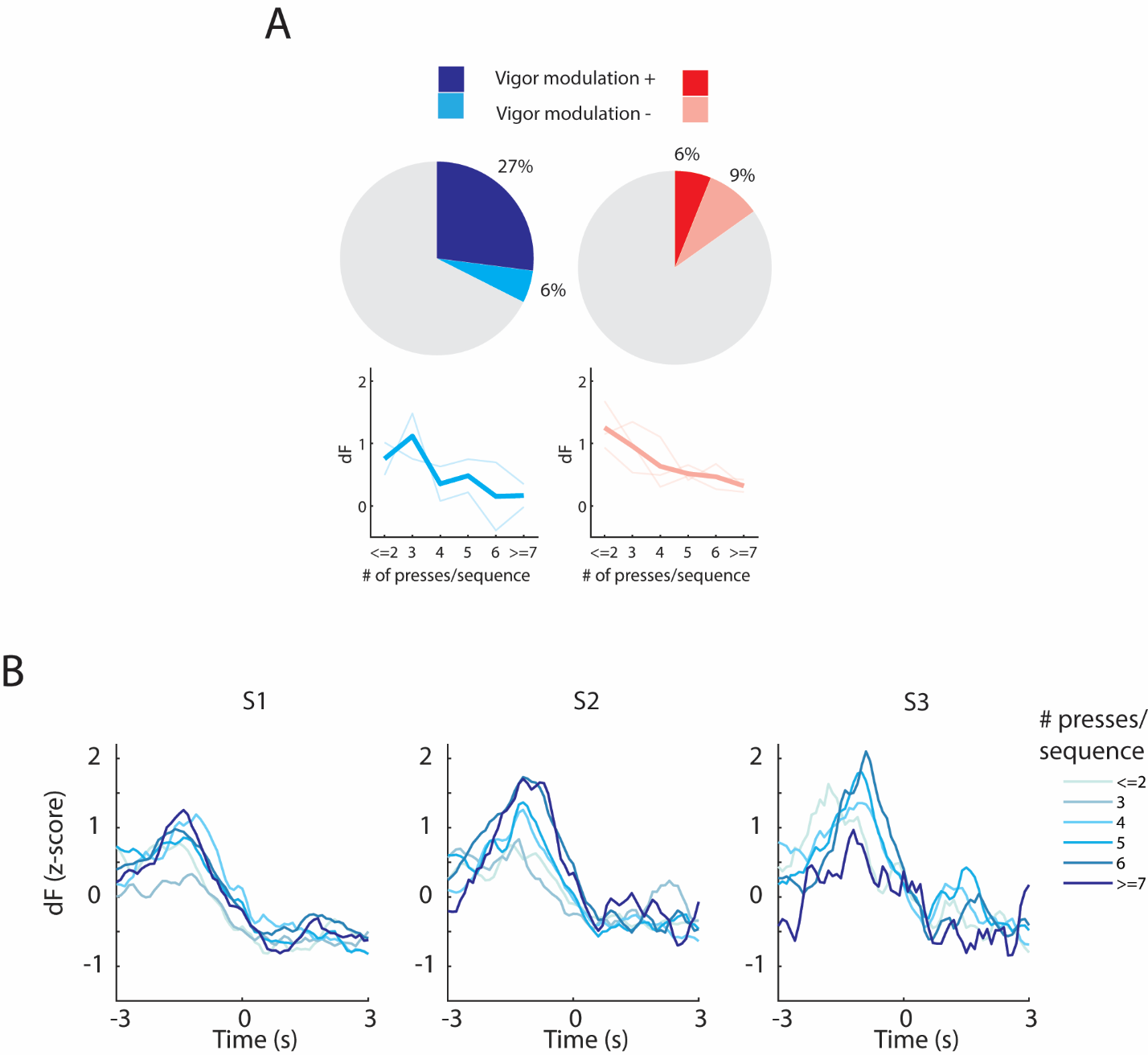

**Fig. S6.**

**Transient SNc activity before the first lever press encodes the vigor of contralateral movement sequences.** **A)** Number of negatively modulated neurons in the ipsi and contralateral conditions. **B)** Activity of one example neuron (the one presented in Figure 3B) in Session 1, 2 and 3 sorted by number of presses/sequence in the contralateral sequence performance.

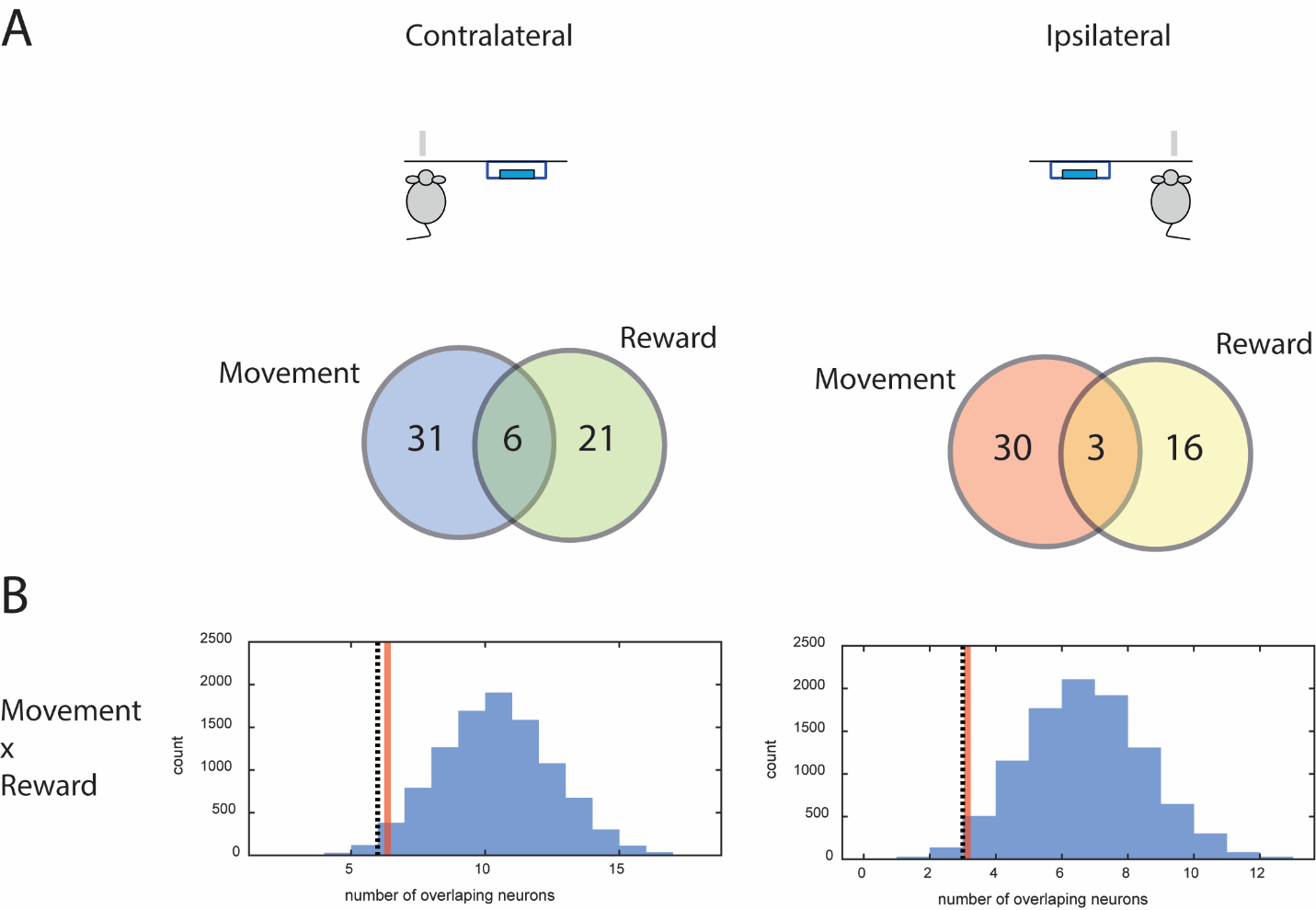

**Fig. S7.**

**Overlap between reward-modulated neurons and movement initiation neurons is minimal and lower than expected by random allocation**. **A)** Overlap between Movement and Reward Neurons. **B)** Monte Carlo simulations (10,000 samples) were used to generate a distribution of the number of overlapping neurons for first press and reward, assuming random assignment. Red line denote the lower margin of the one-sided 95% confidence interval of this simulation (upper limit is +∞). Dashed line represents the number of overlapping neurons found in our experiments.

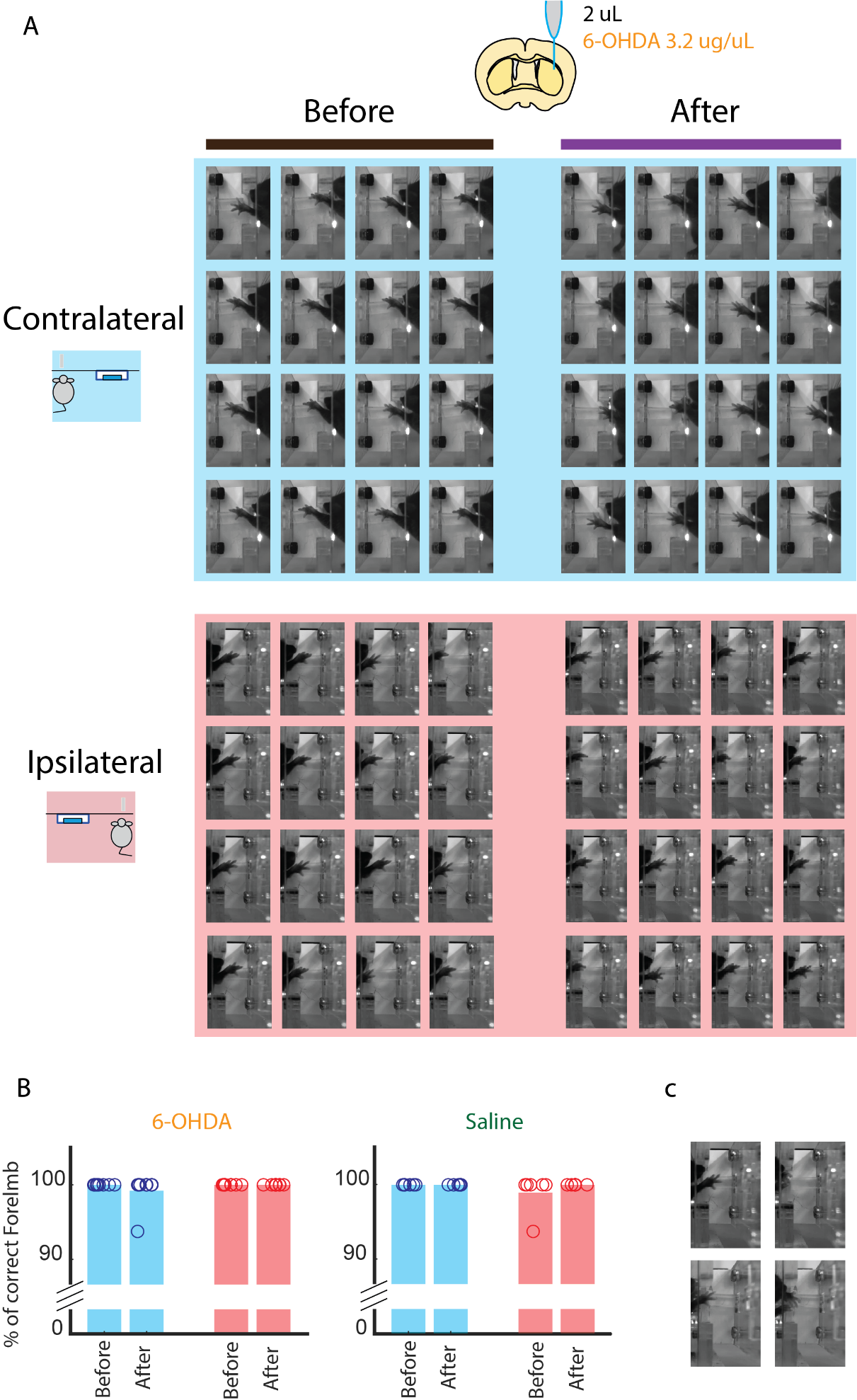

**Fig. S8.**

**Mice still solve the task using the forelimb contralateral to 6-OHDA lesion**. **A)** Still images from randomly selected lever presses of an example 6-OHDA treated mouse. Sixteen (16) randomly selected lever lever presses are used as an example in the 4 conditions (ipsi/contralateral forelimb and before/after 6-OHDA lesion). The mouse used the experiment-intended forelimb in all situations. **B)** For each condition and each animal 16 stills from randomly selected lever presses were visually inspected to assess the use of the correct. Two mice performed 1/16 (6.25%) of the inspected lever presses/condition with the incorrect forelimb (one 6-OHDA treated mouse in the contralateral forelimb after lesion and one saline treated mouse in the ipsilateral forelimb before). Percent of usage of the correct forelimb did not significantly changed after 6-OHDA lesion (Contra: 100% +- 0% to 99.21% +- 0.78; Ipsi: 100% +- 0 to 100% +- 0) or saline injection (Contra: 100% +- 0% to 100% +- 0; Ipsi: 98.96% +- 0.90 to 100% +- 0). **C)** The two situations with incorrect forelimb used are represented on the right images. An example of the same mouse using the correct forelimb is provided for comparison (left).

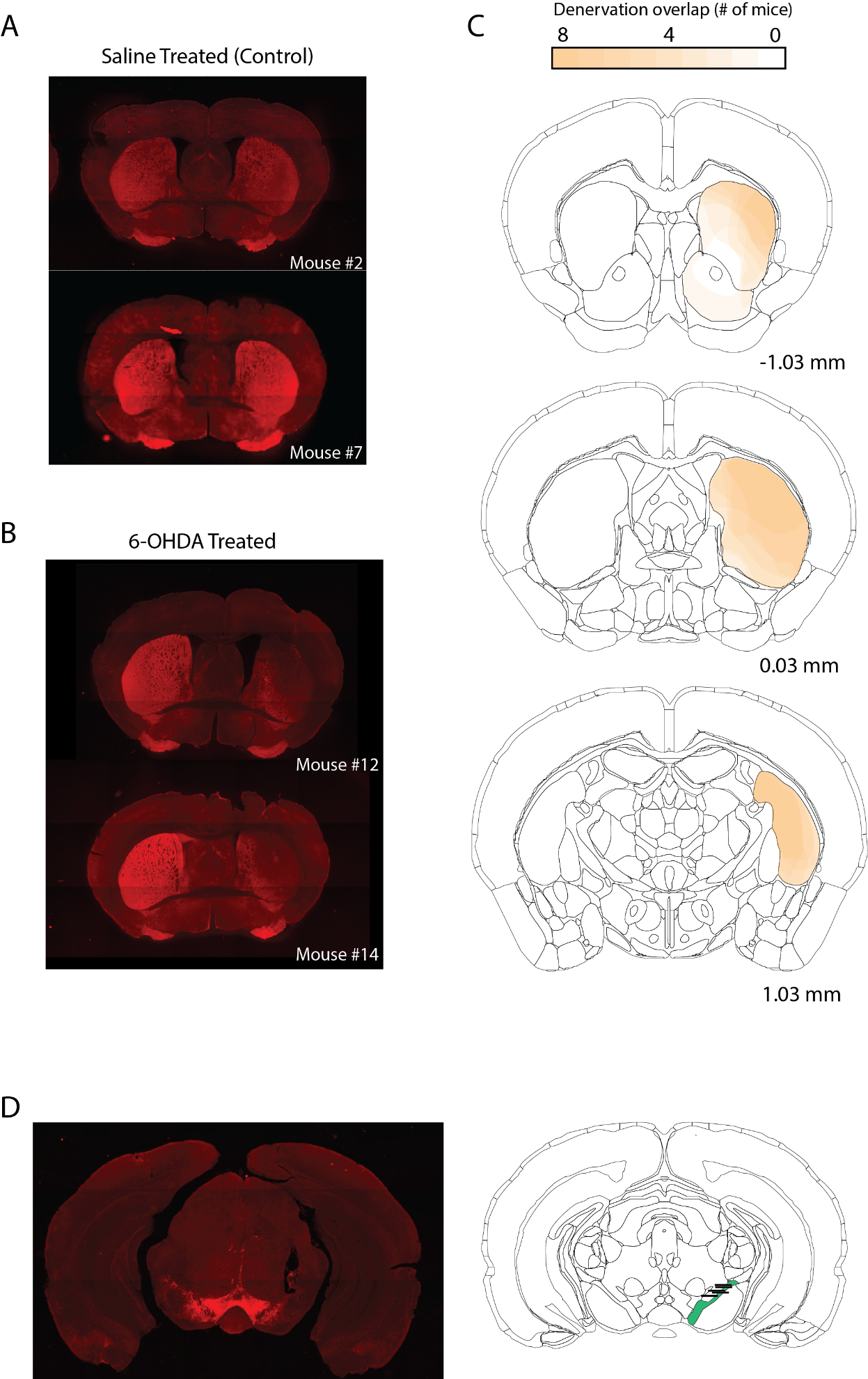

**Fig. S9.**

**Histological assessment of lens placement and 6-OHDA lesions**. **A)** Thyrosine Hidroxylase staining of 2 example mice treated with 2 uL of intrastriatal saline (control group of Figure 5). **B)** Thyrosine Hidroxylase staining of 2 example mice treated with 2 uL (3.2 ug/uL) of intrastriatal 6-OHDA (lesioned group in Figure 5). **C)** Schematics representing overlap of 6-OHDA lesion across the 8 mice. Images were rotated as necessary to display the overlap of lesions in the same hemisphere (part of the animals were treated in the left striatum and the other part in the right). **D)** Thyrosine Hidroxylase staining of one of the mice with lens implant (left). Location of implanted lenses in 5 mice. Bars denote approximate position of lens tip. Lens trajectory was not possible to properly assess in one mouse due to severe tissue damage during perfusion.

Table S1.

Detailed statistical analysis of main figures

| Fig | Sample size (n) | Statistical tests | Value |
| --- | --- | --- | --- |
| 1B | Wild-type (n=8) | One-way repeated measures ANOVA | Learning: **** F (18, 126) = 4,536 p<0.0001 |
|  |  |  | Performance: F (30, 210) = 1,174 p=0.2544 |
| 1C | Wild-type (n=8) | Mixed Effects | Learning: **** F (18, 109) = 7,526, p<0.0001  Learning Inset: **** F (18, 109) = 8.856, p<0.0001 |
|  |  |  | Performance: F (30, 200) = 1,283, p=0.1605  Performance Inset: F (30, 200) = 1,359, p=0.1121 |
| 1D | Wild-type (n=8) | Mixed Effect | L: F (18, 109) = 0.7912 , p=0.7065 |
|  |  |  | P: * F (30, 200) = 1.642, p=0.0247 |
| 1E | Wild-type (n=8) | Mixed Effect | L: **** F (18, 109) = 10.52 p<0.0001 |
|  |  |  | P: F (30, 200) = 1,191, p=0.2383 |
|  |  | One sample t-test (vs. 4) | t7=0.5169, p=0.6212 |
| 1I | Wild-type (n=8) | Mixed Effect | L: ** F (18, 108) = 2,134, p=0.0089 |
|  |  |  | P: F (30,207) = 1.015, p=0.4507 |
|  |  | One sample t-test (vs. 1/3) | t7=0.3619, p=0.7281 |
| 1J | Wild-type (n=8) | Mixed Effect | L: F (18, 108) = 0.8126, p=0.6819 |
|  |  |  | P: * F (30, 200) = 1.736, p=0.0142 |
| 1K | Wild-type (n=8) | Mixed Effect | L: **** F (18, 109) = 3,250. p<0.0001 |
|  |  |  | P: F (30, 200) = 1.309, p=0.1423 |
| 2J | % of Positively modulated neurons (n=6) | Paired t-test | t=0.357, df=5, p=0.7356 |
| 2K | Positively modulated neurons:  Contra: n=37  Ipsi: n=33 | Unpaired t-test | t=2.014, df=68, *p=0.0480 |
| 3A | Positively modulated neurons. Contra: n=37: Ipsi: n=33 | Mixed effects analysis  Fixed effect  Test for Linear trend  Slope of linear fit | Contra:  F (5, 168) = 4.235, p=0.0012  **** F (1, 168) = 20.65, p<0.0001  Slope (95% CI): 0.0639 (0.0361 – 0.0916)  Ipsi:  F (5, 151) = 1.470, p=0.2030  F (1, 151) = 0.4521 p=0.5023  Slope (95% CI): -0.0086 (-0.0347 – 0.0170) |
| 3C | Vigor modulated neurons. | Fisher Exact test | * Contra: n=10/37: Ipsi: n=2/33; p=0.0266 |
| 3F | ROIs correlation  n=114 | Paired t-test | **** t=14,66, df=113, p<0.0001 |
| 3G | Positively modulated neurons. Contra: n=37: Ipsi: n=33 | Mixed effects analysis  Fixed effect  Test for Linear trend  Slope of linear fit | Contra:  F (5, 175) = 6.207, p<0.0001  **** F (1, 175) = 24.69, p<0.0001  Slope (95% CI): 0.0607 (0.0366 – 0.0848)  Ipsi:  F (5, 157) = 1.038, p=0.2913  F (1, 157) = 1.202 p=0.2470  Slope (95% CI): 0.0112 (-0.0099 – 0.0345) |
| 3H | Vigor modulated neurons. | Fisher Exact test | * Contra: n=9/37: Ipsi: n=1/33; p=0.0151 |
| 4C | % of Positively modulated neurons (n=6) | Paired t-test | t=0.2668, df=5, p=0.8003 |
| 4F Left | % of Positively modulated neurons (n=6) | Paired t-test | t=0.1452. df=5, p=0.8902 |
| 4F Right | Positively modulated neurons  Contra: n=27  Ipsi: n=19 | Unpaired t-test | t=0.974, df=44, p=0.3355 |
| 4G Left | % of Positively modulated neurons (n=6) | Paired t-test | t=0.1156, df=5, p=0.9125 |
| 4G Right | Positively modulated neurons  Contra: n=12  Ipsi: n=22 | Unpaired t-test | t=2.723, df=32, p=0.0104 |
| 5C left | Presses/sequence (n=8) | Repeated measures two-way ANOVA | **** Time: F (1, 7) = 68.90 P<0.0001  Forelimb: F (1, 7) = 4.704 p=0.0667  * Time x Forelimb: F (1, 7) = 11,11 p=0.0125  Sidak’s multiple comparison test  *** Contralateral p=0.0005  Ipsilateral p=0.1380 |
| 5C right | Normalized presses/sequence (n=8) | Paired t-test | ** t=3,759, df=7 p=0.0071 |
|  |  | One sample t-test | **** Contralateral: t=11.07, df=7 p<0.0001  Ipsilateral: t=2.281, df=7 p=0.0565 |
| 5D left | Presses/sequence (n=6) | Repeated measures two-way  ANOVA | * Time: F (1, 5) = 7.704 p=0.0391  Forelimb: F (1, 5) = 2.007 p=0.2157  Time x Forelimb: F (1, 5) = 0.01041 p=0.9227 |
| 5D right | Normalized presses/sequence (n=6) | Paired t-test | t=0.4441, df=5, p=0.6755 |
| 5E left | % of Long sequences (n=8) | Repeated measures two-way ANOVA | *** Time: F (1, 7) = 30.12 p=0.0009  Forelimb: F (1, 7) = 2.087 p=0.1918  *** Time x Forelimb: F (1, 7) = 32.45 p=0.0007  Sidak’s multiple comparison test  *** Contralateral p=0.0003  Ipsilateral p=0.9725 |
| 5E right | Change in long sequences (n=8) | Paired t-test | ** t=4,126, df=7 p=0.0044 |
|  |  | One sample t-test | **** Contralateral: t=15,46, df=7 p<0.0001  Ipsilateral: t=0.2100. df=7 p=0.8397 |
| 5F left | % of Long sequences (n=6) | Repeated measures two-way ANOVA | Time: F (1, 5) = 4.911 p=0.0775  Forelimb: F (1, 5) = 0.8281 p=0.4046  Time x Forelimb: F (1, 5) = 0.003268 p=0.9566 |
| 5F right | Change in long sequences (n=6) | Paired t-test | t=0.5099, df=5 p=0.6319 |

Table S2.

Detailed statistical analysis of Figure S1.

| Fig | Sample size (n) | Statistical tests | Value |
| --- | --- | --- | --- |
| A | Wild-type (n=8) | Repeated Measures ANOVA | **** L: F (18, 126) = 4.254, p<0.0001  **P: F (30, 210) = 2.088, p=0.0.0025 |
| B | Wild-type (n=8) | Repeated Measures ANOVA | ** L: F (18, 126) = 2.569, p=0.0012  ** P: F (30, 210) = 2.006 p=0.0025 |
| C | Wild-type (n=8) | Mixed Effect | * L: F (18, 109) = 1.928, p=0.0205  P: F (30, 200) = 1.317, p=0.1371 |
| D | Wild-type (n=8) | Mixed Effect | L: F (18, 109) = 1.692, p=0.0513  P: F (30, 200) = 1.082, p=0.3610 |

Table S3.

Detailed statistical analysis referring to Figure S3. Repeated-measures, 2 way ANOVA. Groups are composed of 6 mice, performing on 2 sides across 3 sessions.

|  | Session | Side | Session x Side |
| --- | --- | --- | --- |
| A: Number of Presses | F(2,10) = 1.330  p=0.3075 | F(1,5)=0.2092  p=0.6666 | F(2,10)=1.290  p=0.3175 |
| B: % of non-isolated Presses | F(2,10) = 1.032  p=0.3914 | F(1,5)=0.03333  p=0.8623 | F(2,10)=1.094  p=0.3719 |
| B: % of Seq > 1 (inset) | F(2,10) = 1.219  p=0.3358 | F(1,5)=0.3852  p=0.5620 | F(2,10)=0.8604  p=0.4521 |
| C: Number of Rewards | F(2,10) = 6.091  p=0.0186 | F(1,5)=0.05358  p=0.8261 | F(2,10)=0.2880  p=0.7558 |
| D: % of Rewarded Sequences | F(2,10) = 2.179  p=0.1640 | F(1,5)=0.3660  p=0.5716 | F(2,10)=0.07044  p=0.9324 |
| E: Presses/Sequence | F(2,10) = 1.985  p=0.1880 | F(1,5)=2.919  p=0.1483 | F(2,10)=0.4165  p=0.6703 |
| G: IPI | F(2,10) = 2.009  p=0.1847 | F(1,5)=0.3257  p=0.5929 | F(2,10)=0.4805  p=0.6320 |
| I: IPI fano factor | F(2,10) = 1.513  p=0.2667 | F(1,5)=0.004525  p=0.9490 | F(2,10)=0.9085  p=0.4340 |
| J: Number of Sequences/minute | F(2,10) = 3.992  p=0.0532 | F(1,5)=0.08217  p=0.7859 | F(2,10)=0.1292  p=0.8803 |
| K: Number of Rewards/minute | F(2,10) = 3.912  p=0.0556 | F(1,5)=0.2349  p=0.6484 | F(2,10)=0.001934  p=0.9981 |
| L: Maximum Press Speed | F(2,10) = 4.075  p=0.0508 | F(1,5)=0.03228  p=0.8645 | F(2,10)=0.6672  p=0.5346 |

Movie S1.

Mice learn to perform sequences of Lever Pressing with only one forepaw. Simultaneous display of the top and side camera on real time. The animal develops a stereotyped behavior going from the magazine to the lever, pressing a few times and then returning to consume the reinforce.

Movie S2.

Mice learn to perform sequences of Lever Pressing with only one forepaw. Low-speed video (x0.2) of press sequences present in Movie S1.

Movie S3.

Example mouse performing the task with the paw ipsilateral to 6-OHDA injection. Left represents before and right represents after injection.

Movie S4.

Example mouse (same animal as in Movie S3) performing the task with the paw contralateral to 6-OHDA injection. Left represents before and right represents after injection.
